# Sperm Whales Demographics in the Gulf of Alaska and Bering Sea/Aleutian Islands: An Overlooked Female Habitat

**DOI:** 10.1101/2023.04.16.537097

**Authors:** Natalie Posdaljian, Alba Solsona-Berga, John A. Hildebrand, Caroline Soderstjerna, Sean M. Wiggins, Kieran Lenssen, Simone Baumann-Pickering

**Author notes:** Corresponding Author (RC).

## Abstract

Sperm whales exhibit sexual dimorphism and sex-specific latitudinal segregation. Females and their young form social groups and are usually found in temperate and tropical latitudes, while males forage at higher latitudes. Historical whaling data and rare sightings of social groups in high latitude regions of the North Pacific, such as the Gulf of Alaska (GOA) and Bering Sea/Aleutian Islands (BSAI), suggest a more nuanced distribution than previously understood. Sperm whales are the most sighted and recorded cetacean in marine mammal surveys in these regions but capturing their demographic composition and habitat use has proven challenging. This study detects sperm whale presence using passive acoustic data from seven sites in the GOA and BSAI from 2010 to 2019. Differences in click characteristics between males and females (i.e., inter-click and inter-pulse interval) was used as a proxy for animal size/sex to derive time series of animal detections. Generalized additive models with generalized estimation equations demonstrate how spatiotemporal patterns differ between the sexes. Social groups were present at all recording sites with the largest relative proportion at two seamount sites in the GOA and an island site in the BSAI. We found that the seasonal patterns of presence varied for the sexes and between the sites. Male presence was highest in the summer and lowest in the winter, conversely, social group peak presence was in the winter for the BSAI and in the spring for the GOA region, with the lowest presence in the summer months. This study demonstrates that social groups are not restricted to lower latitudes and capture their present-day habitat use in the North Pacific. It highlights that sperm whale distribution is more complex than accounted for in management protocol and underscores the need for improved understanding of sperm whale demographic composition to better understand the impacts of increasing anthropogenic threats, particularly climate change.

## Introduction

Male and female sperm whales are sexually dimorphic and the sexes have differences in behavior and habitat preference that result in differences in their distribution and seasonality (1–3). Females and their dependent young form social groups and are known to inhabit temperate and tropical latitudes (2,4). As males mature, they leave their social group and travel to higher latitudes, where they form bachelor groups as juveniles and are mostly solitary as they mature sexually (1,2,4). The males are thought to make periodic migrations to lower latitude breeding grounds once they are sexually mature (1,2,4). Recognizing these demographic differences, sperm whales are managed within the North Pacific stock by the NOAA National Marine Fisheries Service as a single demographic group (5) consisting of only adult males (6).

In the North Pacific, particularly in the Gulf of Alaska (GOA) and Bering Sea/Aleutian Islands (BSAI) regions, most sperm whale distribution data come from a combination of historical whaling data and visual surveys. Social groups were reported in whaling data in the North Pacific as far north as 50°N in the summer (7,8) several records of sperm whales of both sexes overwintering in the western Aleutians (8–14). Estimates for female sperm whale catches range from 6% of total catch above 50°N (8) to 80% in the western Aleutians, western Bering Sea, and the USSR defined Gulf of Alaska (12). More recent surveys report a sighting of a group of females and immature sperm whales in the Central Aleutians (11) and a group of eleven mixed-sex individuals, including one calf in the summer off the continental slope southwest of Kodiak Island (15). This historical and contemporary evidence demonstrates that social groups are not restricted to temperate and tropical latitudes and that their distribution is more complex than currently represented in management assessments.

Sperm whales are deep-diving cetaceans that spend more than 70% of their time in foraging dive cycles (16). The high proportion of time spent at depth makes them difficult to study using typical visual line-transect surveys, but they are excellent candidates for Passive Acoustic Monitoring (PAM) due to their high-amplitude and easily identifiable echolocation signals (17). Three acoustic studies have documented the presence of sperm whales in the GOA (18–20). Additional recordings from more sites with longer time series would allow for characterization of the spatiotemporal patterns of these animals.

Differences between male and female body size is linked to differences in sperm whale click characteristics (21–23). Sperm whales produce broadband echolocation clicks in the 8 Hz to 20 kHz band, with a distinct spectral shape and a peak frequency at about 10 kHz (24). Male sperm whale echolocation clicks have high source levels (236 dB re 1 μPa at 1 m; Møhl *et al.* 2000) resulting in their detection over long distances (9-90 km; Madsen *et al.* 2002; Barlow & Taylor 2005; Mathias *et al.* 2013). Sperm whale echolocation clicks have a multipulse structure (28), and the time between these pulses is called the Inter-Pulse Interval (IPI). The IPI is a result of the time taken for the click to reflect multiple times between air sacs at opposite ends of the spermaceti organ and to exit the rostrum in several subsequent pulses (24,29). Since the length of the spermaceti organ or the rostrum of the animal is about one-third of the total body length (30), stereo photogrammetry measurements of body length and the speed of sound in the spermaceti organ allow for the derivation of two equations (based on two different populations) that relate IPI measurements to body length (21,22). Several studies have used manual and automatic extraction methods to estimate the acoustic length from IPIs recorded by acoustic tags and single sensor instruments (31,32). Average IPI values range from 2-9 ms which translates to an acoustic body length estimate of 7.7 to 17.8 m. A key application of these studies is to differentiate male and female animals based on their IPI and inferred body size.

Due to source directionality, most recorded sperm whale clicks do not display a clear multi-pulse structure and tend to have complex pulse trains (32–34). This limitation results in sparse information about demographic composition since the number of IPI measurements that are possible from acoustic recordings is limited. An alternative approach is to use the Inter-Click Interval (ICI), which is the time between pulse trains, as a proxy for sperm whale body size and sex, particularly for large-scale acoustic monitoring where clicking bouts can last for hours (23). Adult males and females also have different ICIs, with males clicking every ~1s and females clicking every 0.5s (35), which is like other odontocetes that display a relationship between ICI and body size (36).

In this study, we used acoustic recordings of sperm whale echolocation clicks and the differences in their ICI to derive spatiotemporal patterns for male and female sperm whales at five sites in the GOA and two sites in the BSAI spanning the years 2010 to 2019. These data were investigated on an hourly and daily level to understand temporal and spatial habitat use. Generalized additive models (GAMs) with generalized estimation equations (GEEs) were used to evaluate significant spatiotemporal patterns for males and females and compared to available literature, including historical whaling data. Additionally, we used Generalized Linear Models (GLMs) to explain the relationship between sperm whale presence and drivers of presence like small- and large-scale climate variability. This study provides a baseline for sperm whale demographic presence and builds on spatiotemporal patterns described in a region experiencing environmental change. The demographic complexities revealed in this study suggest the need to re-evaluate management of the North Pacific stock, which currently only accounts for adult male presence. Demographic specific responses to climate change should be accounted for to develop the most effective plans for conservation and protection of this species.

## Methods

### Data Collection

Passive acoustic recordings were collected at seven sites, two along the BSAI and five in the GOA, between June 2010 and September 2019 (Fig 1; Table 1). Each site had from one to ten deployments which resulted in ~12 years of cumulative recordings between all sites. Individual site temporal coverage varied due to project goals, recorder battery life, data storage space, and duty cycle regimes as detailed below. The sites were in moderate water depths of 780 m to 1200 m (Table 1). We used High-frequency Acoustic Recording Packages (HARPs; 37) with a sampling rate of 200 kHz which can detect the high-frequency echolocation clicks of odontocetes, including but not limited to, sperm whales.

**Fig 1.**
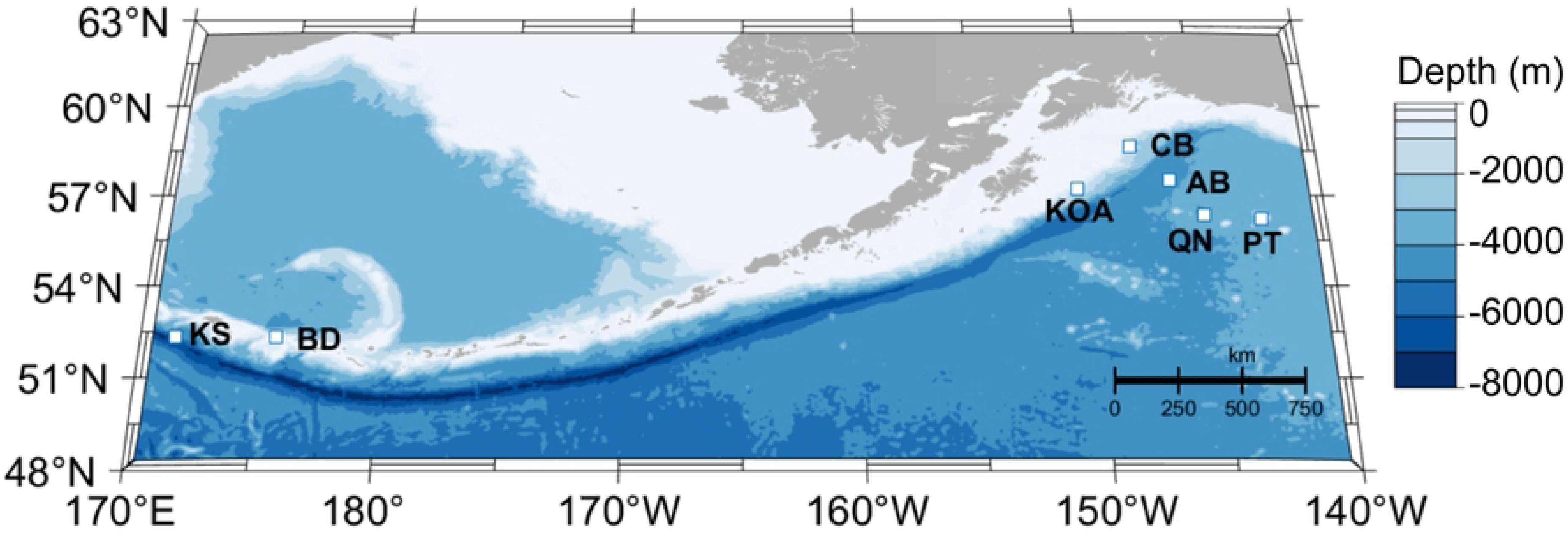
Recording locations (square markers with site abbreviations) in the GOA and BSAI regions. Bathymetry represented by blue color scale in meters.

**Table 1.** Summary of recording effort in the GOA and BSAI regions from 2010 to 2019. Each row represents an individual deployment. Recording effort includes region, site name (abbreviation), latitude, longitude, depth, recording dates, and total number of recording days for each deployment with the site total bolded in the final row. Deployments marked with an asterisk (*) have a duty cycle: the second continental slope (CB) deployment had a 10-minute recording duration every 12 minutes and the second Buldir Island (BD) deployment had a five-minute recording duration every ten minutes.

| Region | Site | Latitude (N) | Longitude (W) | Depth (m) | Recording Dates (MM/DD/YY) | No. of Recording Days |
| --- | --- | --- | --- | --- | --- | --- |
| Gulf of Alaska (GOA) | Continental Slope (CB) | 58° 38.74' | 148° 04.13' | 1000 | 07/13/11 – 02/19/12 | 221 |
|  |  | 58° 40.28' | 148° 01.25' | 900 | 05/03/12 – 02/21/13* | 294 |
|  |  | 58° 40.41' | 148° 00.55' | 877 | 06/06/13 – 09/05/13 | 91 |
|  |  | 58° 40.31' | 148° 01.31' | 858 | 09/05/13 – 04/28/14 | 235 |
|  |  | 58° 40.26' | 148° 01.46' | 914 | 04/29/14 – 09/09/14 | 133 |
|  |  | 58° 40.25' | 148° 01.46' | 900 | 09/09/14 – 05/02/15 | 235 |
|  |  | 58° 39.32' | 148° 05.48' | 929 | 05/01/15 – 09/06/15 | 128 |
|  |  | 58° 40.26' | 148° 01.45' | 874 | 04/30/17 – 09/12/17 | 135 |
|  |  | 58° 40.22' | 148° 01.62' | 900 | 09/14/17 – 06/16/18 | 275 |
|  |  | 58° 40.18' | 148° 01.57' | 972 | 04/25/19 – 09/27/19 | 155 |
|  |  |  |  |  |  | **1902** |
|  | Pratt Seamount (PT) | 56° 14.61' | 142° 45.44' | 989 | 09/09/12 - 06/10/13 | 275 |
|  |  | 56° 14.64' | 142° 45.43' | 987 | 06/11/13 - 08/20/13 | 70 |
|  |  | 56° 14.58' | 142° 45.41' | 988 | 09/03/13 - 03/21/14 | 199 |
|  |  | 56° 14.60' | 142° 45.46' | 987 | 04/30/14 - 09/10/14 | 134 |
|  |  |  |  |  |  | **678** |
|  | Quinn Seamount (QN) | 56° 20.36' | 145° 11.24' | 930 | 09/11/13 - 04/16/14 | 217 |
|  |  | 56° 20.48' | 145° 10.99' | 900 | 09/10/14 - 05/2/15 | 234 |
|  |  | 56° 20.44' | 145° 11.11' | 994 | 05/02/15 - 08/18/15 | 109 |
|  |  | 56° 20.48' | 145° 10.99' | 964 | 04/30/17 - 09/14/17 | 138 |
|  |  |  |  |  |  | **698** |
|  | Abyssal Deep (AB) | 57° 30.82' | 146° 30.05' | 1200 | 04/28/17 - 09/14/17 | **139** |
|  | Kodiak Island (KOA) | 57° 13.44' | 150° 31.70' | 1000 | 04/24/19-09/27/19 | **157** |
| Bering Sea and Aleutian Islands (BSAI) | Buldir (BD) | 52° 38.00' | 175° 37.99' | 783 | 08/27/10 - 05/26/11 | 272 |
|  |  | 52° 04.56' | 175° 38.39' | 777 | 05/31/11 – 08/11/11* | 438 |
|  |  |  |  |  |  | **712** |
|  | Kiska (KS) | 52° 19.01' | 178° 31.24' | 1092 | 06/03/10 – 08/26/10 | **84** |

### Detecting Sperm Whales

Sperm whale echolocation clicks were detected using the multi-step approach described in Solsona-Berga *et al.* 2020 (appendix). These clicks have multiple pulses (28), 2-9 ms apart, depending upon the size of the animal (29). As a result, the detector had a lockout for clicks separated by less than 30 ms to avoid multiple detections of a single click. Band passing the data (5-95 kHz) minimized the effects of low-frequency noise from vessels, weather, or instrument self-noise on detections, but allowed for detection of the echolocation clicks of toothed whales. The Power Spectral Density (PSD) of detected signals was calculated with the *Pwelch* method (MATLAB, 39) using 4 ms of the waveform and a 512-point Hann window with 50% overlap (40). Instrument specific full-system transfer functions were applied to account for the hydrophone sensor response, signal conditioning electronics, and analog-to-digital conversion. To provide a consistent detection threshold, only clicks exceeding peak-to-peak (pp) sound pressure level (RL) of at least 125 dBpp re 1 µPa were analyzed. This threshold was chosen to eliminate noise signals and the echolocation clicks of other odontocetes, while retaining sperm whale clicks.

Sperm whale echolocation clicks can be confused with the impulsive signals from ship propeller cavitation. An automated classifier developed by Solsona-Berga *et al.* 2020 (appendix) was used to exclude periods of ship passages. The classifier identified potential ship passages from long-term spectral averages (LTSA), which are long duration spectrograms (37). Further averaging was calculated as Average Power Spectral Densities (APSD) per 2-hour blocks over low (1-5 kHz), medium (5-10 kHz), and high (10-50 kHz) frequency bands with 100 Hz bins and 50% overlap. Using received sound levels, transient ship passage signals were separated from odontocete echolocation clicks and weather events. A trained analyst manually reviewed identified ship passages using the MATLAB-based custom software program *Triton* (37). Ship passage times were removed from further analysis and considered time periods with no effort.

Instrument self-noise and the echolocation clicks of other odontocetes were also removed to reduce the number of false positive detections. A classifier using spectral click shape was implemented, taking advantage of a sperm whale click’s distinct low-frequency spectral shape to remove dissimilar clicks by delphinid and beaked whales, which typically have higher frequencies (38). The remaining acoustic encounters containing putative sperm whale echolocation clicks were manually reviewed with *DetEdit*, a custom, MATLAB-based graphical user interface (GUI) software program used to view, evaluate, and edit automatic detections (38).

### Click Characteristics as a Proxy for Demographics

Histograms of ICI provide a visualization that can be used to indicate sperm whale size and sex (23). A plot of concatenated histograms, referred to as ICIgrams, was annotated and categorized for each time period at each site. Examples of the ICIgram GUI can be found in Solsona-Berga *et al.* (2022). We used three ICI groups to correspond to three size classes (Fig 2, bottom panels), as per Solsona-Berga *et al.* (2022). Detections with a modal ICI of 0.6 s or less were presumed to be females and their young, hereinafter referred to as Social Groups. Detections with a modal ICI of 0.8 s and greater were presumed to be adult males, hereinafter referred to as Adult Males. The detections with a modal ICI between the Social Groups and Adult Males (< 0.6 s and > 0.8 s) could contain large females or juvenile males, hereinafter referred to as Mid-Size.

**Fig 2.**
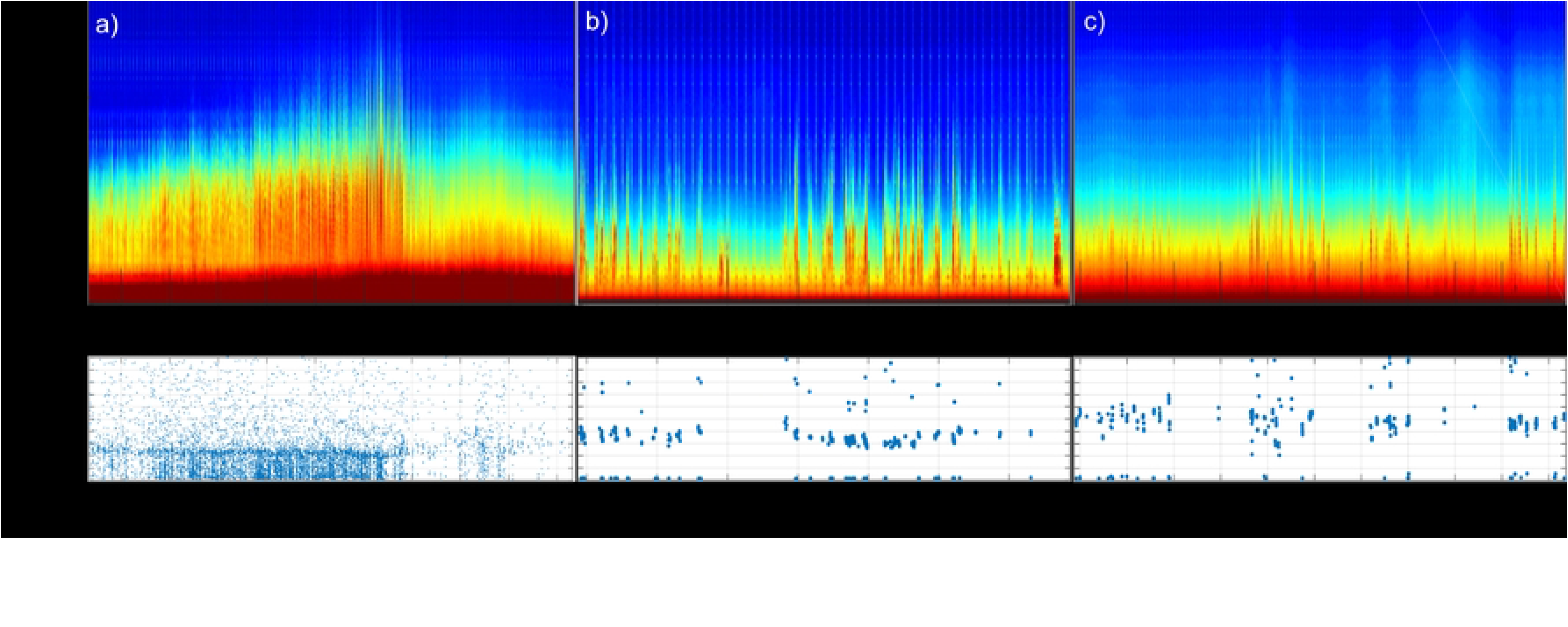
Sperm whale echolocation clicks in long-term spectral average (LTSA; top panel) with their time between detections (ICI; bottom panel). The panels represent three different modal ICIs: a) 0.5 s, b) 0.7 s, and c) 1.0 s. The size of the points on the first panel (a) were minimized for ease of visualization of the modal ICI.

The ICIgram method was originally developed for sperm whales in the Gulf of Mexico (23), where the population is known to have small body size and consist mostly of females, their calves, and immature animals (41,42). To compare how effectively the ICIgram method can be used to categorize the size/sex of sperm whales in the GOA/BSAI, length estimates using IPI from individual animals were matched with the size/sex classification using the ICIgram method.

IPIs were extracted using the Cachalot Automatic Body Length Estimator (CABLE) of Beslin *et al.* (2018). This tool estimates the body length of sperm whales by compiling and clustering their IPI distributions. To avoid including the same animal more than once, only unique IPI values were retained in the final analysis. The length of the whale was estimated using two equations derived from regression analysis of IPI measurements and photogrammetrically estimated body lengths. The Gordon (1991) equation was developed based on measurements from eleven sperm whales off Sri Lanka and was applied to animals less than 11 m in length (31). The Growcott *et al.* (2011) equation was developed based on measurements from 33 large male sperm whales off Kaikoura, New Zealand and was applied to animals greater than 11 m in length (31).

### Click Detection Processing

Sperm whale click detections were binned into 5-minute intervals. The mean daily presence per week was calculated by summing the number of 5-minute bins with detections for each size class and for each site. Since not all sperm whale clicks were categorized into a size class, a time series of unclassified clicks was also included for each site. The ratio of hourly as well as daily presence for each size class was calculated and displayed with Venn diagrams to show the overlap of the classes at each site. Finally, these data were grouped into one-hour bins for statistical modeling, as described in the next section. The one-hour bins were chosen as a compromise to maintain data granularity while ensuring at least 30 minutes of recording effort in each one-hour bin for the two duty-cycled deployments.

### Statistical Modeling

Generalized Additive Models (GAMs; 43) combined with Generalized Estimating Equations (GEEs; 44) were used as a model framework, outlined by Pirotta *et al.* (2011), in the software *R* (46) to test the significance of temporal predictors on sperm whale presence. Patterns were explored for all sperm whales combined, hereinafter referred to as the Inclusive model, and for each of the three size classes, referenced as the Social Group, Mid-Size, and Adult Male models. Models were built for each of these groups for each site with more than 270 days of recording (BD, CB, PT, and QN), for each region (GOA and BSAI), and for an All-Site model. The response variable was binomial presence-absence of sperm whale clicks in one-hour bins (1 = presence and 0 = absence). The explanatory variable *Julian day* was included for all site-specific models while the variable *year* was only included at CB where more than five years of data were available. The region-specific models included *Julian day* and *site* (BD, KS, CB, PT, QN, AB, KOA) as explanatory variables. *Year* was only included in the regional GOA model where more than five years of data were available. Finally, for the All-Site model, *Julian day*, *region* (GOA and BSAI), and *year* were included. The variable *time lost* was originally included as the number of missing 5-minute recording bins in each hour to account for the differences in recording effort due to ship passage exclusions but ultimately removed from final models due to a lack of significance.

Sperm whale encounters lasted for many hours to days at all sites, indicating temporal autocorrelation whereby detections in a single one-hour bin increased the likelihood of detections in adjacent bins. To minimize the impacts of the temporal autocorrelation and to avoid data sub-sampling or using a coarse analysis resolution, the GAMs were combined with GEEs, a method previously used to address autocorrelation in marine mammal presence data (45,47–49). Under this approach, the data are grouped into blocks, within which residuals are allowed to be correlated, while independence is assumed between separate blocks. The *R* correlation function *acf* within the *stats* package (46) was used to determine the time step for blocking. Blocks were defined by the value where the autocorrelation of the residuals of a Generalized Linear Model (GLM) dropped below 0.1. Block sizes varied between 226 - 1249 hours (9 – 52 days) for all 28 models. Although GEEs are considered robust against correlation structure misclassification (44), an autoregressive order 1 (AR-1) covariance structure was used to describe model error given the temporal autocorrelation in the data (45,47,49–52).

The same GLM used to determine block size was also used to assess collinearity of covariates following (53). The *vif* (Variance Inflation Factor) function in the *R* package *car* (54) identified potentially collinear covariates. None of the variables in the GLM model had a VIF value over 2.0 and all variables were retained for further modeling.

Models were built using the function *geeglm* in the *geepack* library (55) in *R*. Variables were treated differently (spline vs. factor) within each model based on the nature of the covariate. Given the long time series at each site and region, *Julian day* was included as a cyclic spline based on a variance-covariance matrix built using the *gam* function in the *mgcv* package in *R* (56) to fit a circular smooth in a GEE framework. Given the small number of years for the time series, *year* was included as a factor to estimate year specific effects. *Site* and *region* were input into models as factors given the categorical nature of both variables.

For models with more than one variable, model selection used the Quasilikelihood under Independence model Criterion (QIC_a_) value, an alternative to Akaike’s Information Criterion for GEE models (57), available through the function *QIC* in the *geepack* library in R (v.1.1-6; 58). Manual backwards stepwise model selection was carried out where the model with the lowest QIC_a_ from the full model (all variables) and a series of models containing all terms but one, was used in the following step (45,48). This selection method continued until removing any of the remaining covariates caused the QIC_a_ to increase. The order of the variables in the final model was determined by which variable, when removed, increased the QIC_a_ the most. A Wald’s Test was conducted on the final model using the function *anova* in the *geepack* library to access the significance of each variable in the model. Any non-significant covariates were removed from the models using backwards stepwise model selection until all p-values of the remaining covariates were greater than 0.05. Partial-fit plots for each variable in the final models were created using the approach of Pirotta *et al.* (2011). The x-axis for *Julian Day* is represented by the months of the year and interpreted as sperm whale occurrence among seasons (winter: December - February, spring: March - May, summer: June – August, fall: September – November).

Goodness of fit for the models was evaluated using the *performance* package in *R* (59). The coefficient of discrimination, also known as Tjur’s R2 (60), was calculated for each model using the function *r2_tjur*. Binned residuals were also used to assess the fit of the models. Binned residual plots were obtained using the function *binnedplot* (61). A good fit was expected to have residuals within the 95% confidence intervals (61).

Generalized Linear Models (GLMs) examined the relationship between sperm whale presence and the El Niño Southern Oscillation’s (ENSO) via the Oceanic Niño Index (ONI), the Pacific Decadal Oscillation (PDO) index, the North Pacific Gyre Oscillation (NPGO) index, and the Marine Heatwave Watch (MHW). The monthly PDO, ONI and NPGO values were extracted using the *rsoi* package in *R* (62) and the MHW forecast was generated using Jacox *et al.* (2022). Hourly binary presence of sperm whales was averaged for each month and divided by the recording effort. To remove seasonality, the timeseries was deseasoned using the functions *stl* and *seasadj* in the *forecast* package in *R* (64,65). Previous studies in the GOA found an 8–12-month lag between ENSO events and sperm whale peak presence (66). And since PDO, ONI, and NPGO are connected to one another (67), 8–12-month lags were tested for these indices as well.

## Results

### Comparison of IPI and ICI

Sperm whale body length estimates were calculated using both their IPI and ICI for 3,047 animals encountered across four sites. An effort was made to account for site, seasonal, and interannual variability. The animal lengths obtained from the IPI were divided into the size/sex classes obtained from the ICIgram and the results visualized using violin plots (Fig 3). These plots reveal clear distinctions between the ICI classes based on the body lengths measured by IPI. The Social Groups class is comprised of small animals with a median length of 10.2 m (n = 2,387) and a moderate interquartile range (IQR) of 1.9 m (the range of the middle 50% of the distribution). The Adult Males class has large animals with median length of 15.7 m (n = 325) and a small IQR of 1.2 m. Whereas the Mid-Size class has median of 13.6 m (n = 335) and a broad range of body lengths with an IQR of 6.6 m. There were outliers within the Social Groups and Adult Males classes, where the length estimates from their IPIs indicated that the ICIgram method may have misclassified the size/sex of the animal. These usually occurred during encounters when more than one size/sex class were echolocating at the same time.

**Fig 3.**
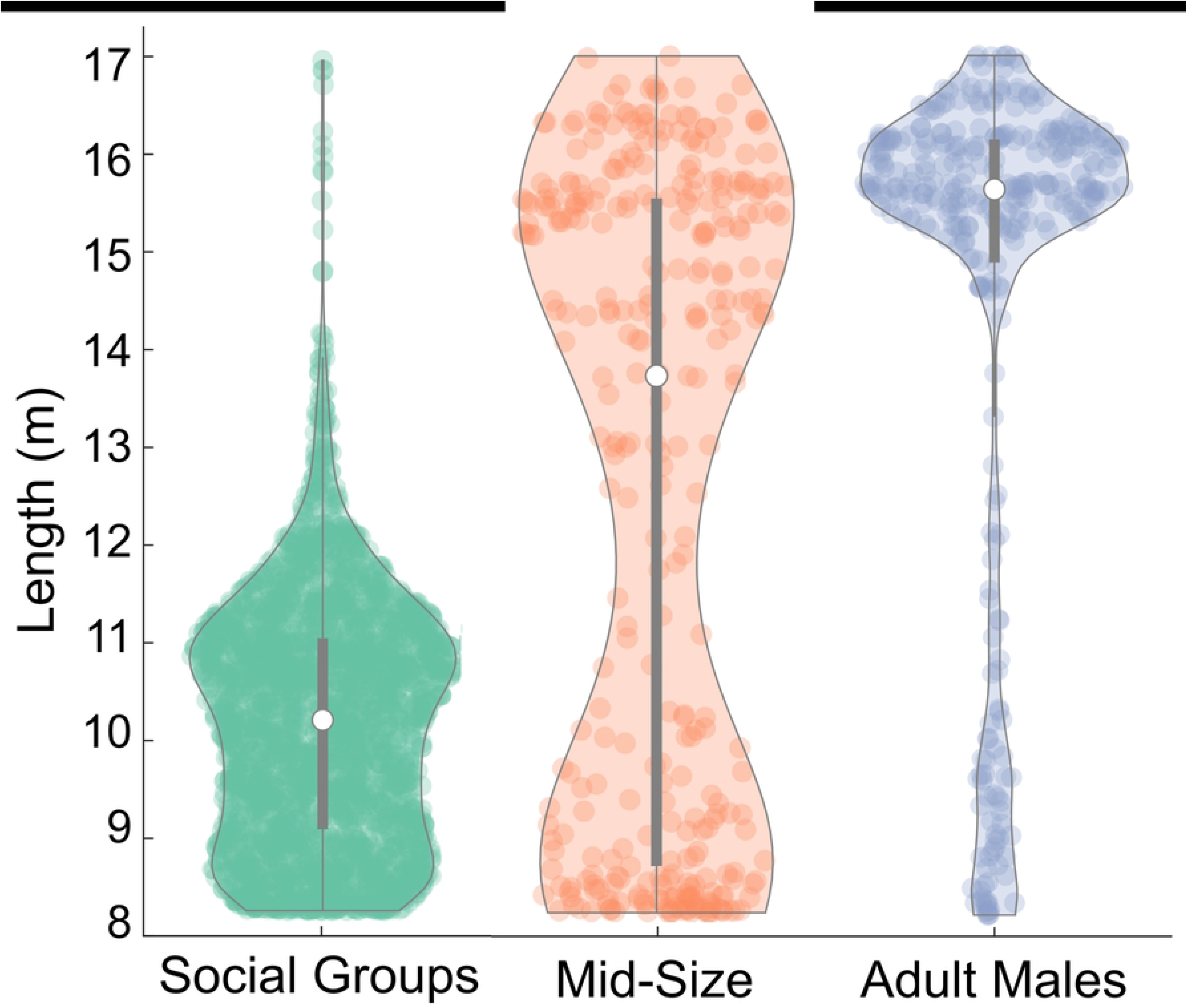
Estimated body length (m) from the IPI categorized into the three ICI size classes (Social Groups, Mid-Size, Adult Males). Median represented by the white dot and the interquartile range by the gray bar.

The distribution of ICI size classes varied between sites (S1 Fig). Averaged across sites, the Social Groups had a mean ICI value of 680 ms, with a range from 600 to 700 ms across sites; the Mid-Size had a mean value of 800 ms, ranging from 750 to 800 ms across sites; and the Adult Males had a mean value of 980 ms ranging from 850 to 1050 ms across sites (S1 Fig).

### Spatial Overlap of Size Classes

All three size/sex classes were detected across all sites, with temporal overlap between classes when observed on both hourly and daily time scales (Fig 4). The highest proportion of overlap at all sites was between Mid-Size and Adult Males. Adult Male and Mid-Size animals were found in the same hourly bin 7% (range 2-16%) and daily bin 36% (range 17-63%) of encounters. Whereas for Social Groups and Mid-Size, they were found together in the same hourly bin only 2% (range 0-5%) and daily bin 8% (range 2-17%) of encounters. Similarly, Adult Males and Social Groups were found together in the same hourly bin 2% (range 0-4%) and daily bin 7% (range 2-20%) of encounters. As expected, encounters with all three size/sex classes were rare with hourly bin overlap of 1% (range 0-2%) and daily bin overlap of 5% (range 0-15%).

**Fig 4.**
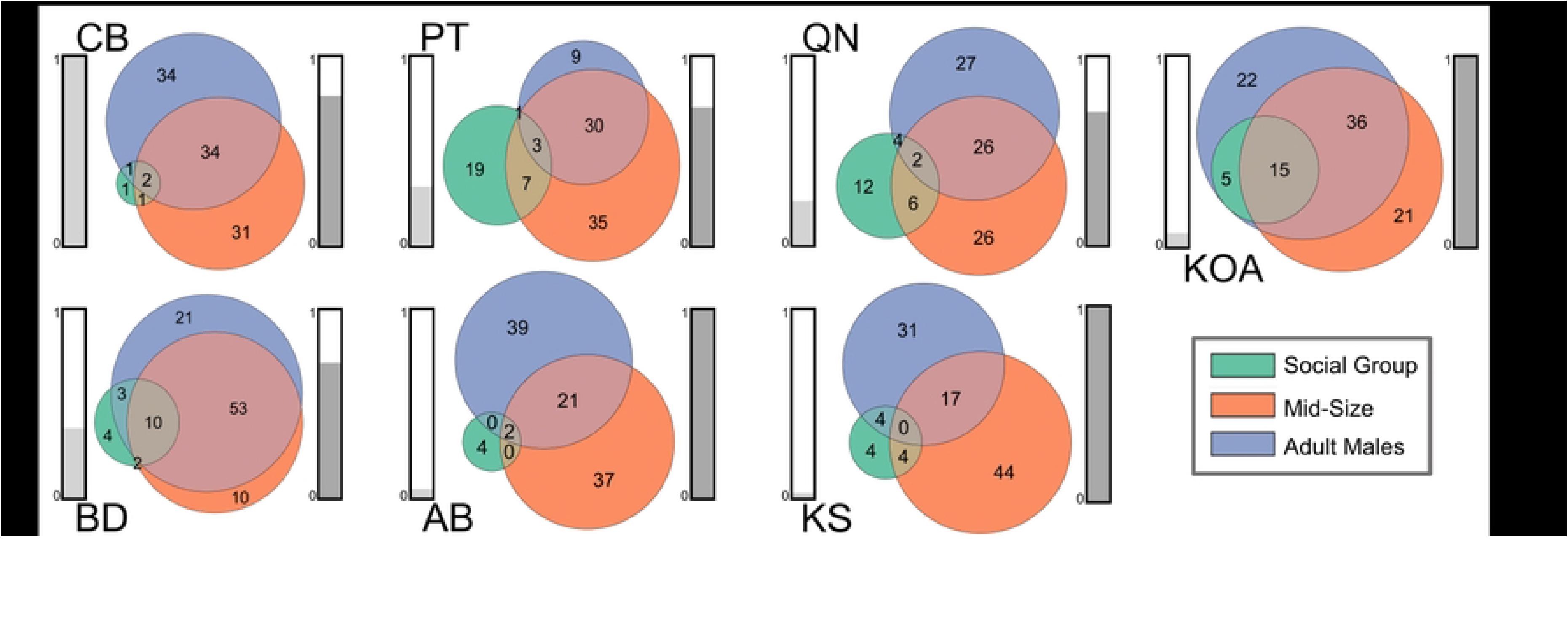
Ratio of hourly (left) and daily (right) presence of each size class at each recording site. Social Groups in green, Mid-Size in orange, and Males in blue. Overlap between groups represents simultaneous presence of those groups in the same hour or day. The bars on the left of each diagram (light grey) represent normalized recording effort at that site. The bars on the right (dark grey) represent normalized sperm whale presence at that site.

At all sites, the proportion of Mid-Size and Adult Male presence was greater than Social Group on the hourly and daily scale (Fig 4). Sites CB, AB, and KOA, along the continental slope and deepwater of the GOA, had the smallest proportion of Social Group presence, while the seamount sites PT and QN had the highest. The proportion of Mid-Size and Adult Males were similar at all sites except for PT where the proportion of Mid-Size presence was the largest (Fig 4).

### Presence by Site

Sperm whales of all size/sex classes were detected at every site and presence was reported as the mean daily presence (min and hr) per week, herein after referred as daily presence. AB, KS, and KOA had the highest normalized daily sperm whale presence and the lowest normalized recording effort (Fig 5, Fig 6). All three sites only captured 19, 12, and 22 weeks respectively, during the spring and summer when sperm whale presence was usually the highest. CB had the next highest normalized daily sperm whale presence and the highest normalized recording effort (> 5 years), followed by PT, BD, and QN (Fig 5, Fig 7).

**Fig 5.**
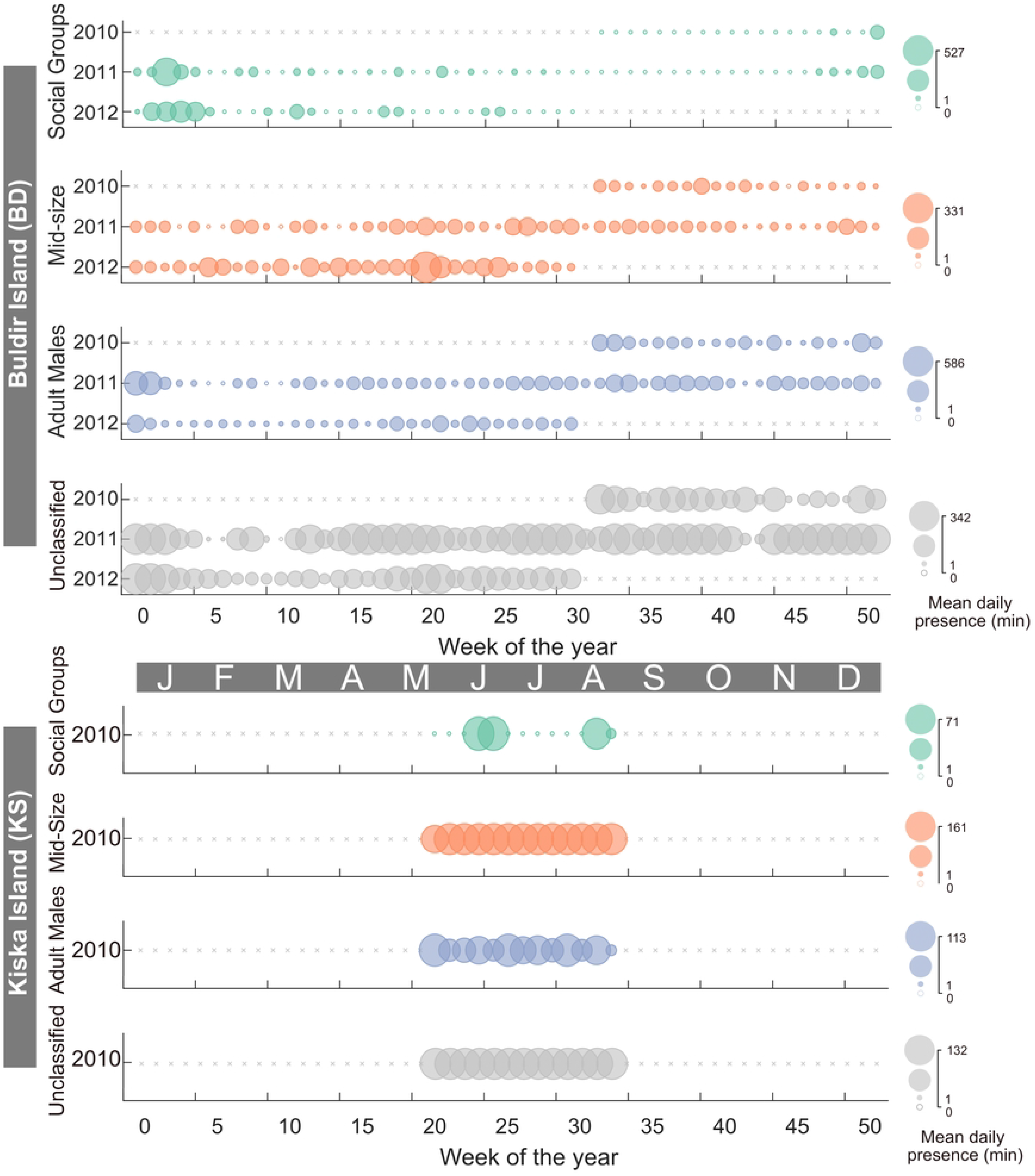
Sperm whale presence at the Buldir Island (BD) and Kiska Island (KS) sites. Each row represents a year. The color of the bubble represents the size class; Social Groups by green, Mid-Size by orange, Adult Males by blue, and unidentified clicks in grey. The size of the bubble is the mean daily presence in minutes represented with a scale on the right. Grey ‘x’ symbols represent no recording effort.

**Fig 6.**
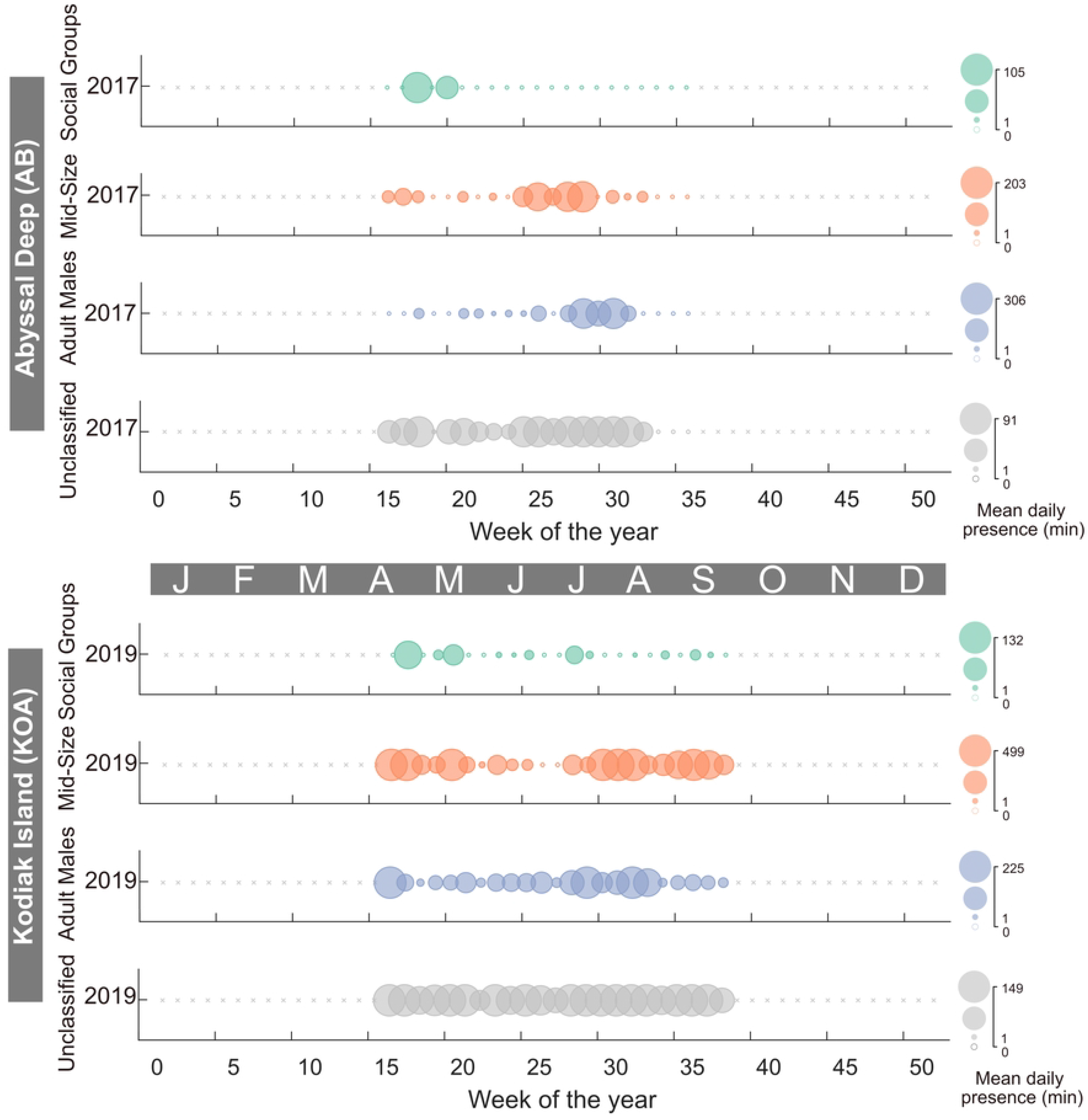
Sperm whale presence at the Abyssal Deep (AB) and Kodiak Island (KOA) sites. Colors and symbols as per Fig 5.

**Fig 7.**
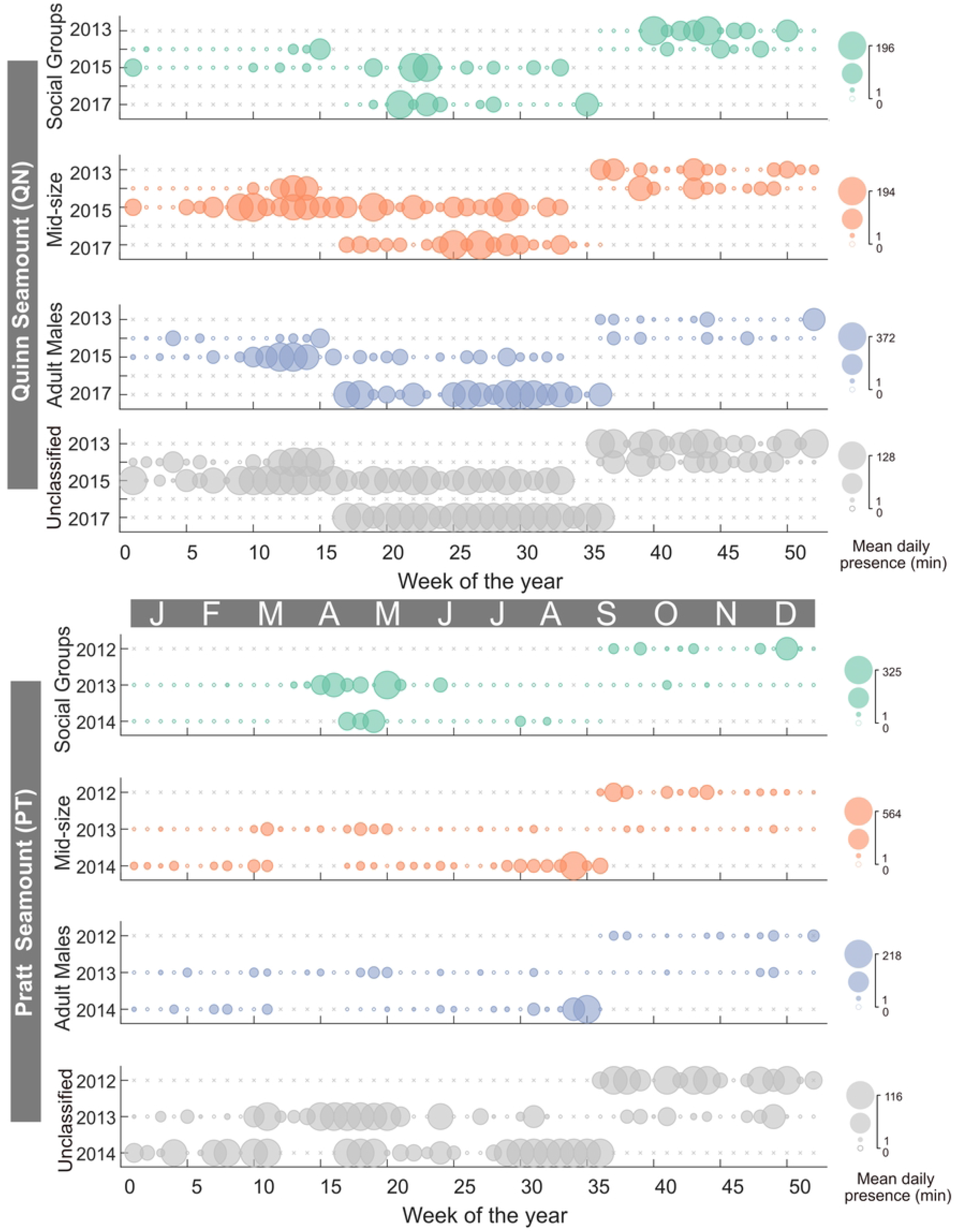
Sperm whale presence at the Quinn Seamount (QN) and Pratt Seamount (PT) sites. Colors and symbols as per Fig 5.

Sperm whales were present almost every week during the nearly two years of recording at the Buldir Island (BD) site (Fig 5). Social Groups were almost exclusively present during the winter months between 2010 and 2012 with a maximum daily presence of 527 min (8.8 h) (Fig 5). Mid-Size were the most consistent size class present with a maximum daily presence of 331 min (5.5 hr) (Fig 5). Adult Males had a maximum daily presence of 586 min (9.7 hr) with the peak in presence seen in January of 2011 (Fig 5). Sperm whales were present every week at the Kiska Island (KS) recording site during the 13-week deployment (Fig 5). Mid-Size and Adult Male presence were higher and more consistent with a maximum daily presence 161 and 113 min (2.7 and 1.9 hr), respectively (Fig 5). Social Groups were present for 4 weeks and had a maximum daily presence of 71 min (1.2 hr) (Fig 5).

In the GOA, the continental slope (CB) site had the highest level of sperm whale presence compared to the other recording sites (Fig 8). Presence at CB was dominated by Mid-Size and Adult Males with maximum daily presence of 846 and 882 min (14.1 and 14.7 hr), respectively. Social Groups were present episodically throughout the eight-year recording period with a maximum daily presence of 104 min (1.7 hr). The two seamount sites, Quinn (QN) and Pratt (PT), had a more consistent presence of all size classes throughout the recording period (Fig 7). The maximum daily presence of Social Groups, Mid-Size, and Adult Males at QN were 196, 194, and 372 min (3.3, 3.2, and 6.2 hr), respectively (Fig 7). The maximum daily presence of Social Groups, Mid-Size, and Adult Males at PT were 325, 564, and 218 min (5.4, 9.4, and 3.6 hr), respectively.

**Fig 8.**
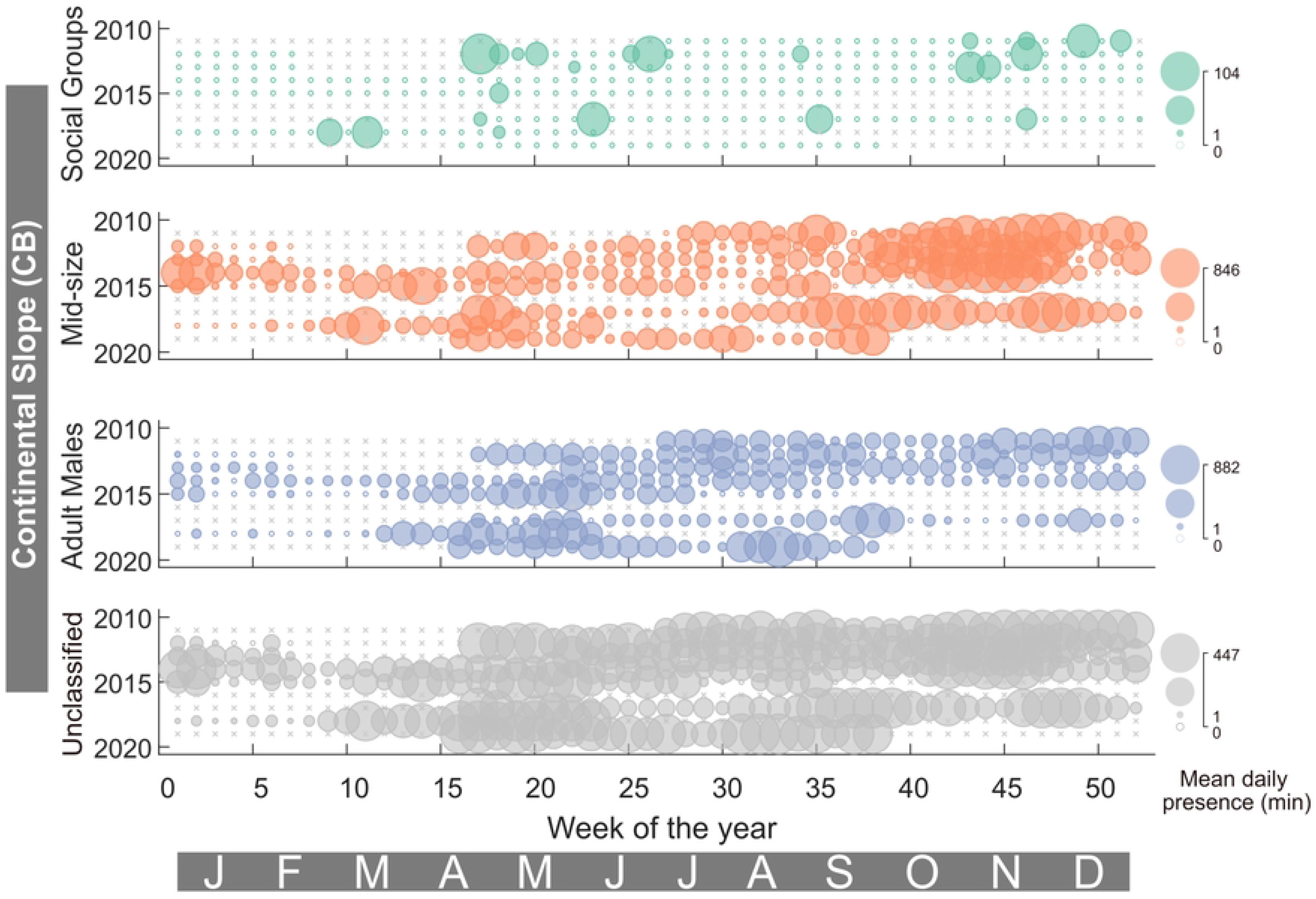
Sperm whale presence for the Continental slope (CB) site. Colors and symbols as per Fig 5.

Sperm whales were present every week of the recording period for the Abyssal Deep (AB) and Kodiak Island (KOA) sites in the GOA. Compared to the other size classes, there was less Social Group presence at AB and KOA with a maximum daily presence of 105 and 132 min (1.8 and 2.2 hr), respectively (Fig 6). There was more presence of Mid-Size and Adult Males at KOA with a maximum daily presence of 499 and 225 min (8.3 and 3.8 hr), respectively (Fig 6). AB had a maximum daily presence of Mid-Size and Males of 203 and 306 min (3.4 and 5.1 hr), respectively (Fig 6).

### Modeling

Sperm whale presence was modeled for all sperm whale classes included together (Inclusive model), and for each of the three size classes independently. Data were used from selected sites with good seasonal coverage (BD, CB, PT, and QN), from all the sites in each region (GOA and BSAI), and from all the sites combined (All-Site).

The average annual percentage of one-hour bins with presence for the Inclusive, Social Group, Mid-Size, and Adult Male models was 98% (8579), 4% (362), 44% (3902), and 32% (2779), respectively (S1 Table). The highest performing models were the Adult Male models with 32 to 50% of Residuals within the 95% confidence intervals. The lowest performing models were the Social Group models with 7 to 25% within the 95% confidence intervals (S1 Table). The models had low Tjur’s R2 values and % of Residuals within the 95% confidence intervals suggesting that the temporal (Julian day and year) and spatial (site) variables included in the models are not good predictors of animal detections.

### Seasonal Patterns

Significant seasonal patterns were found in the majority (26 out of the 28) of models (Table 2). The Inclusive models revealed a seasonal pattern of increased presence in the summer for all GOA sites and fall for the BSAI sites. The patterns revealed by the Inclusive models were like those of the Adult Males at all sites where presence was highest in the summer for GOA and fall for BSAI (Fig 9). The seasonal patterns for Mid-Size and Social Groups were more nuanced and varied from site to site. For the Mid-Size, peak presence was seen in the summer or fall across all sites and regions except QN where peak presence was observed in the spring (Fig 9, S2-S3 Fig). For Social Groups, peak presence was seen in the spring, except for QN and BD where the peak in presence was in the fall and winter, respectively (Fig 9, S2 Fig, S3 Fig). Peak presence of the Social Groups rarely overlapped with those of the Adult Males.

**Fig 9.**
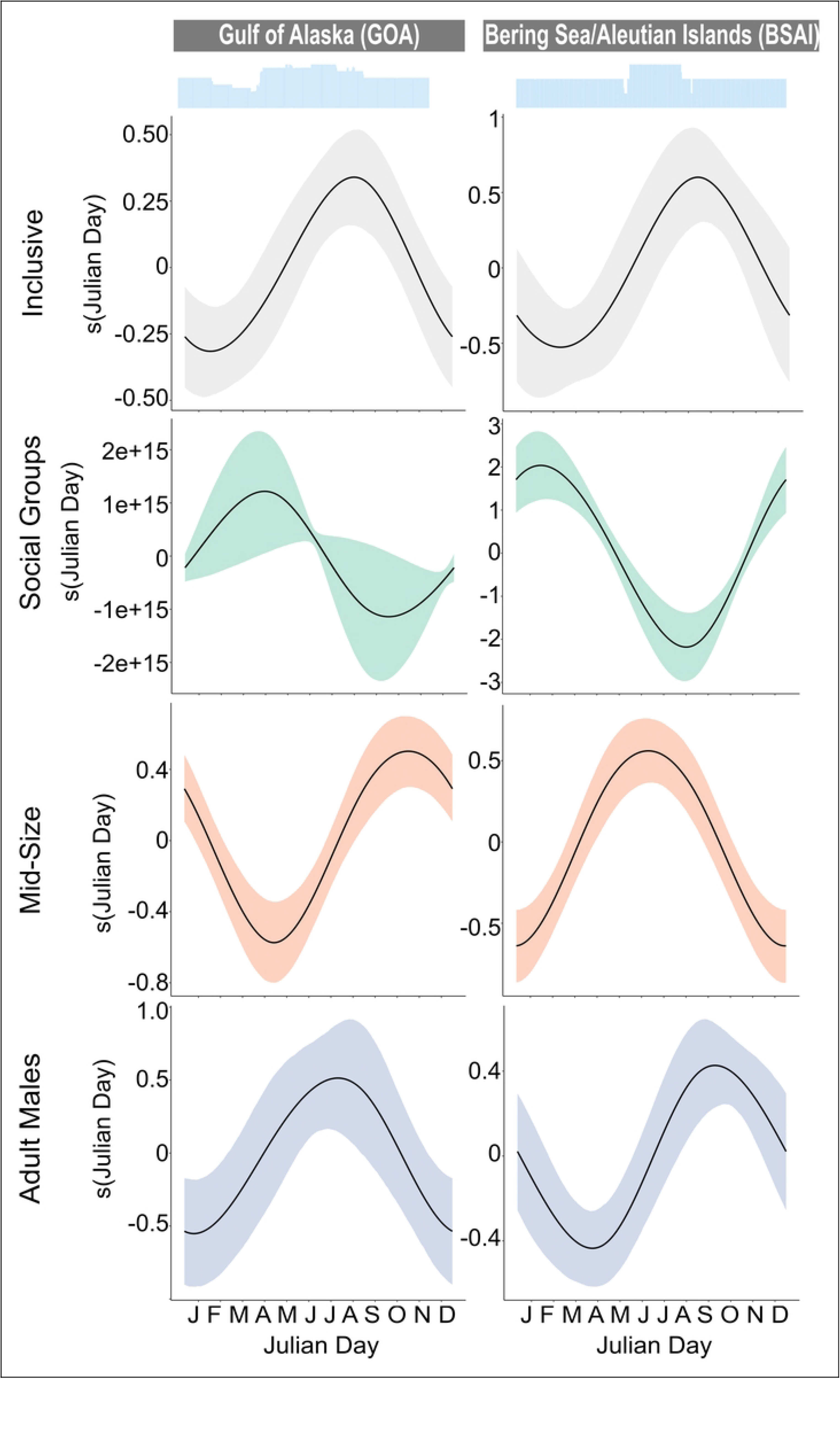
Seasonal plots for the sites in the Gulf of Alaska (left) and the Bering Sea/Aleutian Islands (right). Each row represents outputs from the different size class models for each site: a) Inclusive (grey), b) Social Groups (green), c) Mid-Size (orange), and d) Adult Males (blue). Julian day is represented as months. The blue histograms at the top denote effort. All plots include 95% confidence intervals represented by the grey shading surrounding the smooth.

**Table 2.**
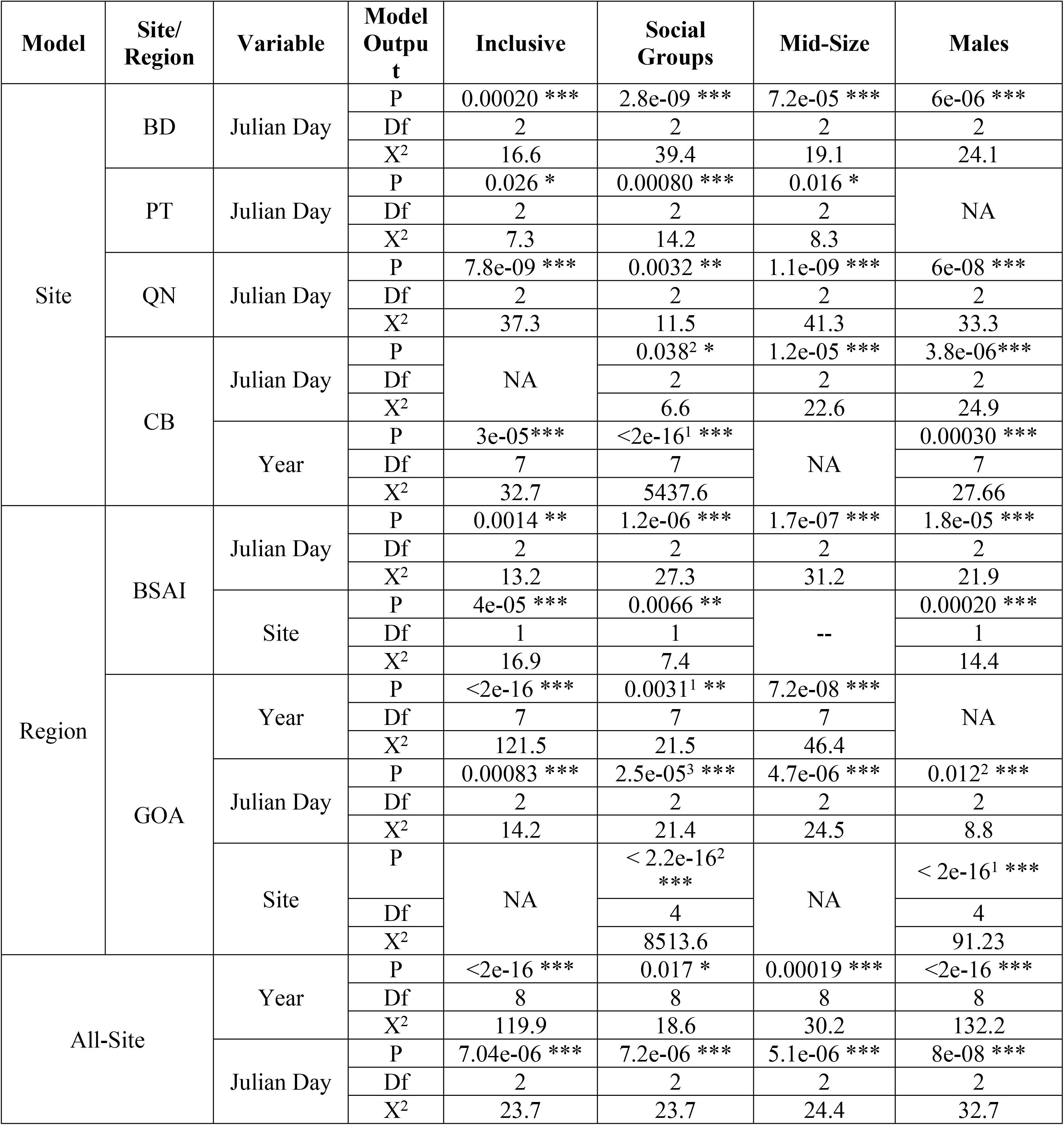
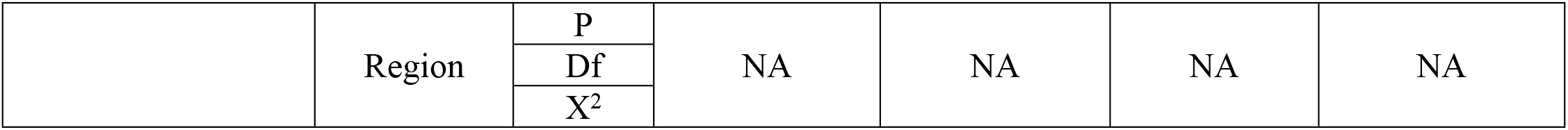
Model summaries for each site, regional, and All-Site models for the Inclusive, Social Groups, Mid-Size, and Adult Male classes. Model summaries include the p-value (P), degrees of freedom (Df), and the Chi-square statistic (X^2^). The significance of the p-value is indicated by the following codes: ‘***’ <0.001, ‘**’ <0.01, and ‘*’ <0.05. If a model had more than one variable, the listed order of the variables represents the order they were input into the model. Models that had different input orders have a subscript for the p-value indicating the order it was input into the model. Covariates that were not retained in the model or not significant are represented with ‘NA’.

### Interannual variability

Interannual variability was only assessed for the CB site, GOA region, and the All-Site models where there was more than five years of data (S3 Fig, Fig 10). At CB, the Inclusive and Adult Male models revealed a decrease in presence every year after 2011 with the lowest presence in 2014, 2015, and 2017, followed by an increase in 2018 and 2019 (S3 Fig). For Social Groups, presence remained steady from year to year except in 2014 and 2019 where there was no Social Group presence whatsoever. Interannual variability was not significant for Mid-Size at CB. In the GOA, the Inclusive and Mid-Size models revealed a decrease in presence every year after 2011 with the lowest presence in 2013 and 2014, followed by an increase every subsequent year (Fig 10). Social Group presence remained steady from year to year with the highest presence in 2011 and a small dip in presence in 2014 and 2015. Interannual variability was not significant for Adult Males in the GOA region. For all seven sites, the Inclusive, Mid-Size, and Adult Male models revealed an increase in presence in 2011, followed by a decrease in presence and a minimum in 2013 with increasing presence in subsequent years. Social Group presence remained consistent from year to year with a dip in presence starting in 2014 and the lowest presence in 2018.

**Fig 10.**
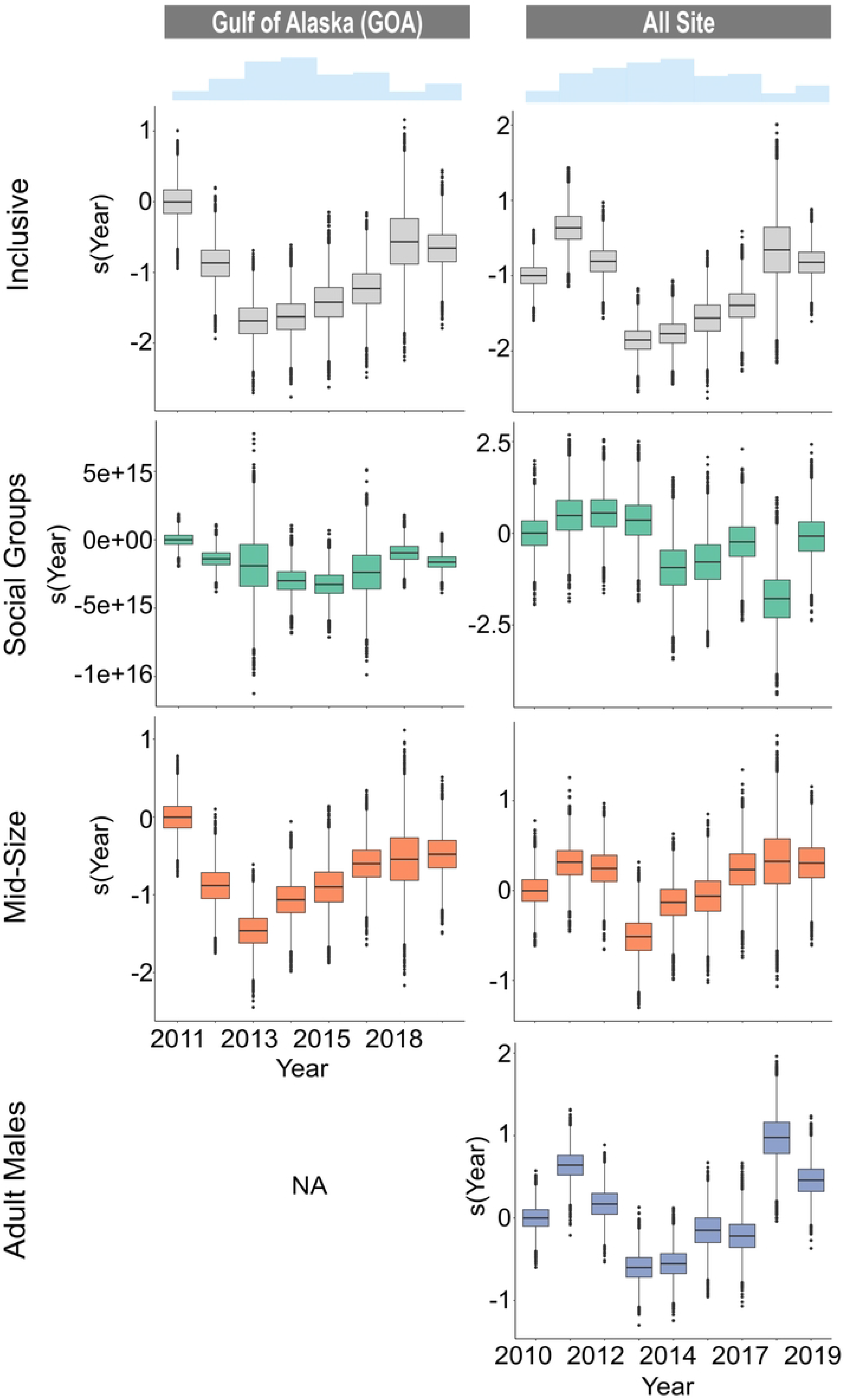
Presence by year for the Gulf of Alaska region (left) and All Sites (including Bering Sea/Aleutian Islands) (right). Each row represents outputs from the different size class models for each site: a) Inclusive (grey), b) Social Groups (green), c) Mid-Size (orange), and d) Adult Males (blue). Year is a categorical variable displayed as box plots with the first level centered on zero. Covariates that were not retained in the model or not significant are represented with ‘NA’.

### Environmental Variability

Sperm whale presence was correlated to the PDO, ONI, and NPGO indices in varying degrees depending on the model. The PDO index with an eight-month lag was significant for all models except for the Mid-Size at CB and Social Groups at GOA (S2 Table). All significant models revealed a negative correlation between the PDO index and sperm whale presence (Fig 11, S9-S10 Fig). The decrease in sperm whale presence in 2013 aligns with the inflection point of the PDO from a cool to warm phase (S6-S8 Fig). The ONI was significant for less than half of the models with a nine-month lag being consistently significant for all models (S2 Table). All significant models revealed a negative correlation between the ONI and sperm whale presence (Fig 11, S9-S10 Fig). The decrease in sperm whale presence in 2013 aligns with the ENSO becoming neutral and is sustained as it transitions to El Nino (warm phase) (S6-S8 Fig). The NPGO index was significant for all models except for the Inclusive at CB, Social Groups at GOA, and all Mid-Size models (S2 Table). Like the PDO index, an eight-month lag was significant for all models except for the Social Groups at CB. All significant models revealed a positive correlation between the NPGO index and sperm whale presence (Fig 11, S9-S10 Fig). The decrease in sperm whale presence in 2013 aligns with the inflection point of the NPGO from a positive to a negative phase (S6-S8 Fig). R^2^ values for all the linear regressions revealed a weak correlation with values ranging between +/−0.27 and +/−0.55 (S2 Table). The MHW forecast was not significant in any of the models (S2 Table) although less sperm whale presence does appear to align with higher MHW probability (S6-S8 Fig).

**Fig 11.**
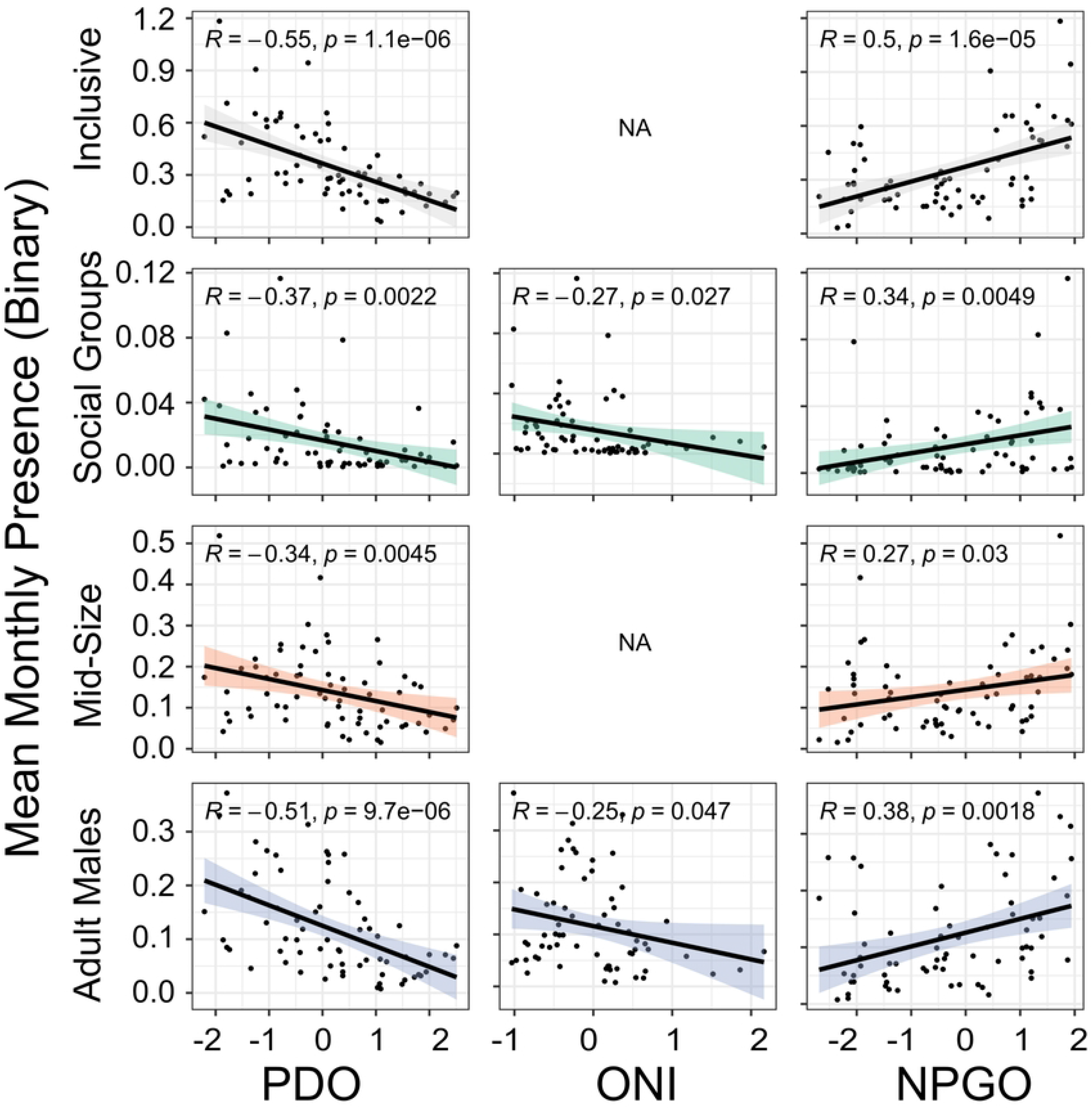
GLM plots displaying the relationship between mean monthly presence of sperm whales for All-Sites and the PDO, ONI, and NPGO index. All PDO and NPGO plots represent an eight-month lag, and all ONI plots represent a nine-month lag. Each row (and color) represents outputs from the different size class models for each variable: Inclusive, Social Groups, Mid-Size, and Adult Males. All plots include 95% confidence intervals represented by the grey shading surrounding the linear regression. The regression formula for each model is displayed in the top left-hand corner. Covariates that were not retained in the model or not significant are represented with ‘NA’.

## Discussion

This study analyzed demographic composition of sperm whales at seven sites in the Gulf of Alaska and Bering Sea/Aleutian Islands over a wide range of years and seasons. Three size/sex classes were identified (Social Groups, Mid-Size and Adult Males) based on their echolocation ICI, supported by examining IPI for individual clicks. The median body length of animals in the Social Group class (10.2 m) is comparable to the average body lengths documented for sperm whale females and immature animals that ranges from 8 to 11 m (1,41,68–71). The median length of the Mid-Size class (13.6 m) is greater than the maximum length for females (12 m) (69–71), suggesting that the Mid-Size group consists of juvenile males. The median length of the Adult Male class (15.7 m) suggests that the males in this study are both physically mature [occurs at a mean length 15.5-15.9 m (72)] and sexually mature [occurs from 9.5 m (30) to 13.8 m (73)]. The seasonal and interannual patterns of the Adult Male and Mid-Size groups show good alignment, further suggesting that the Mid-Size group may consist of juvenile males.

Adult Males were present year-round in the GOA and BSAI, although they were more common in the summer in GOA and fall in BSAI, and less common in the winter and spring. This seasonal occurrence is consistent with what was previously described from acoustic data in the GOA (18) and at Ocean Station PAPA in the southeast GOA (19). Whaling data from the northeastern Pacific also supports this seasonal pattern with an increased mean length of male sperm whales starting between May and June and sustained through September, attributed to the sexual maturity of the animals (3). Male sperm whales in the GOA are also notorious for longline depredation (74) particularly from the sablefish fishery which has its season from mid-March to mid-November (75), aligning with the peak in presence of Adult Males.

The summer peak in presence and winter low in presence is likely associated with long distance movements of males, between lower latitudes where breeding occurs in the winter/spring and higher latitudes, or feeding grounds, for improved foraging opportunities in the summer/fall (2). Although this seasonal trend appears to be migratory in nature, there is little evidence that sperm whales have a predictable pattern and/or route to established breeding areas. Rather, they are described as ‘ocean nomads’ based on Discovery Tags used by whalers, that revealed widespread movements between areas of concentration suggesting that their home ranges can span thousands of kilometers (2,4,8,76). Discovery Tagged animals in southern California and northern Baja California were found in locations ranging from offshore California, Oregon, British Columbia, and the western Gulf of Alaska (8). More recently, studies using satellite tags have corroborated the nomadic behavior of sperm whales. In a study that tagged 10 sperm whales in the GOA, seven stayed within the GOA, one traveled to British Columbia and back, while three traveled south to the Sea of Cortez, Baja, and offshore Mexico through the California Current without stopping and with no synchronized departure (77). Although the three southbound whales did all leave before winter when sperm whale presence was at its lowest, there is no evidence that the animals ‘migrate’ to a specific area outside of their home range. There is also photo-identification evidence from the North Atlantic that sperm whales travel from higher latitudes areas like the Azores, to tropical latitudes like the Gulf of Mexico and Bahamas (78) but no concrete evidence that the animals have a pattern or routine to where and when they travel between presumed higher latitude foraging and lower latitude breeding grounds. Instead, it appears that sperm whales travel in response to the distribution of their often-patchy prey sources (4,79) and are linked with temporary breeding sites with favorable prey conditions driven by the effects of oceanographic conditions. It has also been suggested that sexually mature males don’t breed every year and may choose to remain in higher latitudes some years to feed (80) further complicating their seasonal patterns in and out of their home ranges which can span ocean basins. This is also supported by our acoustic observation of year-round presence of Adult Males in the entire study region. Historical whaling and satellite tag studies provide evidence of highly variable timing and direction of sperm whale movement but incorporating increased observations over longer timescales is necessary to clarify their behavior as nomadic or migratory. Genetic studies reveal that males in the North Pacific have widespread origin and are likely a mix of males from several independent populations in the Pacific (6) further supporting their nomadic nature. If in fact, sperm whales are truly nomadic animals, understanding how they spatiotemporally exploit available resources is important to establishing management and conservation strategies.

Mid-Size animals were also present year-round in the GOA and BSAI, with a slightly offset peak presence from the Adult Males in both regions. The Mid-Size class likely does not undergo long distance movements to breeding grounds since they are sexually immature animals. In the GOA, the peak presence of Mid-Size animals was in the fall or spring months. In the BSAI, peak presence of Mid-Size was in the summer, before the peak presence of Adult Males in the fall, suggesting avoidance of Adult Males by Mid-Size animals. There is evidence of aggression between mature sperm whales based on heavy scarring on their heads (2,81,82). Some juvenile males may avoid an area during peak presence of mature Adult Males to avoid direct competition, although these groups do overlap on an hourly scale in our data, suggesting temporal overlap of habitat use on some level.

Social Groups were present at all seven recording sites but were not present year-round and instead had distinct seasonal patterns that varied from site to site. Social Group presence in the winter months between 2010 and 2012 at site BD is consistent with the 2008 sighting of a group of females and immature animals in the Central Aleutians in winter (11). That sighting was considered rare since only males had been observed in ten years of summer sighting surveys previously conducted in the BSAI region (11). There is also historic whaling evidence that female sperm whales have overwintered in the western Aleutians (8–14). The continued return of Social Groups to this region in the winter, when productivity is generally lower, could be a sign of site fidelity. Although sperm whales have been described as ‘ocean nomads’, there is evidence from females in the Eastern Caribbean (83,84), North Atlantic (85), western Mediterranean (86), and males in the GOA (77) that site fidelity is a factor in their habitat choice. The presence of Social Groups in certain regions of the BSAI in the winter could be evidence of geographic specializations (84).

In the GOA, Social Group peak presence was in the spring. Seasonal prediction models in the waters of coastal British Columbia (BC) found female sperm whales virtually absent after May (3). The absence of females in the BC model predictions suggests that Social Groups could be traveling further north to the GOA in the spring months. The spring peak was also seen in historical whaling data from the northeastern Pacific where female sperm whales were more often caught from March-May and less often caught from June-September (3). Our current understanding of female sperm whale distribution post-whaling does not include the GOA.

Contemporary presence of Social Groups in the GOA and BSAI could represent a return to pre-whaling distributions of sperm whales. Females were illegally caught in high numbers in the North Pacific, removing a significant portion of the reproductively mature population (12). The impacts of whaling on their population, especially given their social ecology, may have been disproportionately large (8,87). Social Group presence could also represent a change in the distribution of their preferred prey, given how closely sperm whale distribution is linked to squid (79,88). The BD recording site was located on the nutrient-rich northern side of the Aleutian Islands in the Bering Sea which would be a prime location for squid and provide suitable habitat for sperm whales. Presence of Social Groups could also be related to changes in the water temperature. Nishiwaki (1966) hypothesized that Social Group presence in the BSAI was related to water temperatures above 13°C. And in the GOA, ocean heat content (HC) was the most important sperm whale predictor, with a decrease in HC leading to a decrease in animal presence (66). However, from 2010 to 2012 when our instruments were recording in the BSAI, this region experienced a multi-year sequential continuation of colder than normal ocean temperatures (89), likely below the 13°C threshold for Social Groups hypothesized by Nishiwaki (1996).

Year-round presence of sperm whales in the GOA and BSAI, especially through the winter, indicates high winter productivity and sustained prey availability (66,90,91). CB had the highest relative presence of sperm whales, particularly of Mid-Size and Adult Males. This site is located along the continental slope which is popular with males in other regions (3) and this site had a sustained presence of sperm whales, even during years with low overall presence in the GOA (15) likely a result of richer biomass productivity. PT and QN, although relatively offshore, correspond to seamounts which are also important sperm whale habitat in the GOA (15) and several other regions due to their complex seafloor characteristics and water circulation (3,92–94). There is evidence from whaling data that females were generally found farther from shore in the northeastern Pacific (95) potentially explaining the higher proportion of Social Group presence at sites PT and QN. Social Groups also appeared linked to oceanographic features and were present as far north as the western North Pacific Gyre in the western Aleutian Islands, and the Alaska Gyre and Alaska Current in the Gulf of Alaska (76,96). In the BSAI, site KS appeared to have the highest relative presence of Mid-Size and Adult Male sperm whales, however, this site has less recording effort than BD and a summer recording effort bias. Regional preference between GOA and BSAI was not significant for any of the size classes, indicating that the two regions are both equally capable of providing suitable foraging conditions.

There were temporal (hourly) and spatial (daily) overlaps between all groups at almost all sites. Temporal and spatial overlap of Adult Males and Social Groups occurred at all sites except AB and could imply that mating is possible in GOA or BSAI. Currently we understand that males travel to tropical latitudes to breed, but there is evidence from whaling data that sperm whales were mating in temperate latitudes off the coast of British Columbia where large bulls were mostly found associated with female schools in April and May (95,97). There is also evidence that the modal breeding month for sperm whales in the North Pacific was April (98) explaining the peak in presence of Social Groups in GOA in spring. Gregr and Trites (2001) hypothesized that by traveling north into temperate latitudes, Social Groups could improve their encounter rates with more mature males that are ready for breeding. There was more spatial than temporal overlap of Adult Males and Mid-Size, due to the offset peaks in presence of the two groups. Less temporal overlap could indicate habitat partitioning or avoidance of sexually mature Males by juvenile males.

Interannual variability of sperm whale presence is due to several ecological, behavioral, and environmental factors related to prey availability in the region, namely squid, fish, and skates (99–103). The dips and peaks of sperm whale presence interannually in our study are supported by visual surveys and reported squid catches. The peak in presence in 2011 for all size classes in this study, were also observed in southeast GOA (66) and could be correlated to the high squid catches reported that year (104). Density and abundance values from visual surveys in the GOA also support the dip in presence seen in our study in 2013 by Adult Males and Mid-Size, with increasing density and abundance in 2015 (15). These dips and peaks in presence are likely a result of changes in prey distribution and abundance which can be difficult to study.

Instead, researchers often rely on understanding how small- and large-scale drivers of ocean productivity optimize feeding and spawning conditions of their prey which ultimately impacts aggregation (105,106). However, the relationship between prey and their environment in the GOA and BSAI, particularly large-scale climate patterns like the PDO, ENSO, and NPGO, is complex and not well understood. Squid are a highly mobile and adaptable group of marine animals that can be found in a wide range of oceanic conditions driven by prey availability and abundance (zooplankton and forage fish), predator populations (salmon, toothed whales, sablefish, and grenadiers), and changes in habitat quality (104). In the GOA and BSAI, there are 15 species of squid whose abiotic habitat preferences are unknown but are likely related to pelagic conditions and currents throughout the North Pacific over various spatial and temporal scales (104). A large climate shift in the mid-1970s from a cold to warm regime (107) resulted in a southward shift and intensification of the Aleutian Low pressure system and warmer ocean temperatures (108). This led to increased zooplankton biomass and demersal, pelagic fish and cephalopod recruitment and abundance (108–110) while forage fish populations declined (111) impacting piscivorous sea birds and some marine mammal populations (110,112,113). While overall the regime shift appeared to increase cephalopod populations potentially as a result of warmer ocean temperatures that accelerate growth rates (114,115) and increased zooplankton biomass, it also resulted in decreased forage fish populations (squid prey) and increased salmon populations (squid predator) (107)and the species-specific impacts of the shift are not well understood. There is also evidence that warmer ocean temperatures and a shoaling of the Oxygen Minimum Zone in the California Current System are resulting in a northward and offshore expansion of some squid species (116–118) creating an environmental refuge in the GOA and BSAI. So, although squid populations might appear to be increasing in warmer ocean temperatures, this increase could be a result of northward range expansion and the impacts on the endemic species are not understood.

In the southeast GOA, peaks in sperm whale acoustic presence seasonally and interannually were related to higher temperatures, a shallow mixed layer, a weaker Alaska Gyre, and enhanced eddy formation (66). These conditions are also associated with El Niño, (ENSO warm phase), (119,120) which has been shown to be positively correlated with peaks in sperm whale presence up to one year later in the southeast GOA (66). El Niño conditions during our recording effort persisted in the GOA from 2014 to 2016 and 2018 to 2019 with very strong conditions from 2015 to 2016 (121). Moderate to strong La Niña (ENSO cool phase) conditions were seen from 2010 to 2012 and a weak La Niña from 2016 to 2018 (121). In this study, opposite to what was seen in the southeast GOA, higher monthly sperm whale presence was associated with La Niña conditions, or a negative ONI. La Niña conditions in the northeastern Pacific are characterized by decreased ocean temperatures, weaker than normal eddies, deeper mixed layer, increased winter nutrient levels, and a return of summer upwelling (122). It is important to note that although secondary effects of ENSO can be felt in the North pacific, it primarily affects lower latitude climates (123).

Larger scale environmental variability such as the PDO could also influence the presence of sperm whales. Like ENSO, the PDO has a positive, or warm phase, and a negative, or cool phase. In fact, these two climate patterns interact with one another and when PDO is highly positive, El Niño will likely be stronger and when the PDO is highly negative, La Niña will likely be stronger (124). A positive PDO brings environmental conditions that have been connected to increased sperm whale presence in the southeast GOA such as higher temperatures and ocean heat content, a shallow mixed layer, and a weaker Alaska Gyre (107,119,125). Our study found a significant negative correlation between PDO and sperm whale presence of all classes. The GLM models with and without lags displayed a significant negative correlation, implying that the effects of the PDO on sperm whale presence span larger time scales. At the start of our recording effort in 2011, the PDO was in a cool phase until it flipped to a warm phase in 2014 (126). This PDO inflection point is also reflected in the sperm whale presence where several low presence years following the shift are associated with a very positive PDO phase. Although the PDO remains in a warm phase for the remainder of our recording effort, the PDO index does decrease dramatically after 2017, and is associated with a steady increase of sperm whale presence through 2019. It is important to note that while our nine-year time series likely does include an important phase switch of the PDO in 2014, the PDO cycle occurs at approximately 20-to-30-year time intervals (107) and it is unlikely that our data are sufficient to capture the full relationship between sperm whale presence and the PDO.

Related closely to ENSO and the PDO, the NPGO is a climate pattern that is significantly correlated to fluctuations in salinity, nutrients, and chlorophyll-a in the Gulf of Alaska (67). Our study found a positive correlation between sperm whale presence and the NPGO index. Sperm whale presence was higher during the positive NPGO phases which are associated with lower SST, higher salinity, chlorophyll-a and nutrients (67). There was no significant correlation between Mid-Size presence and the NPGO index or ONI, implying that juvenile male sperm whales are less linked to ENSO conditions and the NPGO compared to other classes. It is important to note that while the PDO accurately describes climate patterns north of 38°N, the NPGO is most effective at capturing climate patterns south of 38°N (67) which may explain why there was more significance between PDO and sperm whale presence in this study.

Overall, higher sperm whale presence was related to large-scale environmental variability associated with cooler ocean temperatures, higher salinity, chlorophyll-a, and increased upwelling as seen during La Niña, cool PDO phase, and positive NPGO index. Increased nutrient-rich water and higher productivity from La Niña conditions could sustain higher squid populations, although no direct link has been made in the GOA or BSAI. Findings from this study that is focused on the central GOA and BSAI contradict what was seen at Ocean Station PAPA in the southeast GOA (66). Reasons for this include a difference in recording effort; recording at Ocean Station PAPA occurred for five years (66) while this study encompasses nine years of recording effort. It is also possible that the PDO and NPGO are no longer effective tools for predicting changes in marine environments. A study by Litzow et al. 2020 found that since 1988/1989, the main drivers of the PDO and NPGO have become less active, making these large-scale climate patterns less effective at understanding and predicting marine productivity. There is also evidence that the relationship between the PDO and productivity are non-stationary and phase shifts in the PDO can result in distinct climate states that cannot be directly compared (127).

During the period of 2014 to 2016, the northeastern Pacific experienced an unprecedented marine heatwave, often referred to as “The Blob” (128). The more than 2.5°C increase of the upper 100 m of the ocean (129) led to low chlorophyll concentrations that wreaked havoc on marine ecosystems from California to Alaska (130). Low primary productivity likely resulted in poor foraging conditions revealed by the decrease of Mid-Size and Adult Males and complete absence of Social Groups in the GOA. However, there was no significant relationship between the MHW forecast and sperm whale presence for any of the models or classes. This could be a result of no recording effort in 2016 while the marine heatwave continued resulting in the inability of our data to capture the full effects of the marine heatwave in the GOA. Presence began to slowly increase in 2015 and continued to do so until the end of our recording effort. There appeared to be a large increase in presence of Adult Males and a decrease of Social Groups in 2018 for the All-Site model which is likely a result of recording effort bias since there was only acoustic data from one site that year (CB). Climate models for the North Pacific predict environmental changes that would support higher concentrations of prey and attract top predators like sperm whales in high latitudes (66,117,131). Although sperm whale presence in this study appears to increase at the end of the recording period in 2018 and 2019, recording effort bias and the lack of consecutive years with increased presence prevents drawing conclusions about the GOA serving as a foraging refuge for the whales.

Since the spatiotemporal models in this study were only investigating seasonality, interannual trends, and differences in site and region, low model performance was not surprising. The spatiotemporal models would likely be improved with the inclusion of environmental data that is correlated with sperm whale presence such as ocean heat content, sea surface temperature, vertical stratification (66), chlorophyll-a (93,132), mesoscale features like thermal fronts (133) and eddies (93). This study was also limited by short and/or discontinuous time series at certain sites. Since sperm whales display seasonal patterns in this region, some of the models could be biased by recording effort. Continuation of acoustic monitoring at these sites will allow for more robust time series and potentially improve performance of spatiotemporal models. This is particularly important for the Aleutian Island and two seamount sites where Social Group presence was high, highlighting critical habitat for females and their young in this high latitude region.

## Conclusions

This work highlights the importance of understanding sperm whale spatiotemporal distribution and regional demographics for informing appropriate management and conservation measures. Currently, management of the North Pacific stock of sperm whales does not account for Social Group habitat use and assumes that the region is dominated by juvenile and sexually mature males. This study reveals that Social Group presence in this region is likely overlooked and historical presence of females in whaling data and contemporary ‘rare’ occurrences should not be ignored when determining management practices for this stock. Male and female sperm whales have differences in behavior and ecology that likely translate to demographic specific responses to increasing anthropogenic threats and climate change. Creating a baseline understanding of what Social Group presence looks like in the GOA and BSAI is crucial for monitoring future changes to the demographic composition of the North Pacific stock.

## Supporting Information

**S1 Fig.** The distribution of interclick intervals at each site. Social Groups are in green, Mid-Size in orange, and Adult Males in blue. A kernel smoothing function is represented by the bold line outlining the distributions.

(TIFF)

**S1 Table.** Model evaluation summaries for all site-specific, regional, and All-Site models. The number of one-hour bins with presence are given by the # of Bins. The coefficient of discrimination is given by the Tjur’s R^2^. The percent of residuals within the 95% confidence intervals of binned residual plots are given by the % of Residuals.

(DOCX)

**S2 Fig.** Seasonality plots for the two seamount sites in the GOA (PT and QN) and one of the island sites in the Bering Sea/Aleutian Islands (BD). Each row represents outputs from the different size class models for each site: a) Inclusive (grey), b) Social Groups (green), c) Mid-Size (orange), and d) Adult Males (blue). Julian day is represented as months. The blue histograms at the top denote effort. All plots include 95% confidence intervals represented by the grey shading surrounding the smooth. Covariates that were not retained in the model or not significant are represented with ‘NA’.

(TIFF)

**S3 Fig.** Seasonality plots (left) and presence by year (right) for site CB. Each row represents outputs from the different size class models for each site: a) Inclusive (grey), b) Social Groups (green), c) Mid-Size (orange), and d) Adult Males (blue). Year is a categorical variable displayed as box plots with the first level centered on zero. Covariates that were not retained in the model or not significant are represented with ‘NA’.

(TIFF)

**S4 Fig.** Presence by site for the Bering Sea/Aleutian Islands. Each row represents outputs from the different size class models for each site: a) Inclusive (grey), b) Social Groups (green), c) Mid-Size (orange), and d) Adult Males (blue). Site is a categorical variable displayed as box plots with the first level centered on zero. Covariates that were not retained in the model or not significant are represented with ‘NA’.

(TIFF)

**S5 Fig.** Presence by site for the Gulf of Alaska. Each row represents outputs from the different size class models for each site: a) Inclusive (grey), b) Social Groups (green), c) Mid-Size (orange), and d) Adult Males (blue). Site is a categorical variable displayed as box plots with the first level centered on zero. Covariates that were not retained in the model or not significant are represented with ‘NA’.

(TIFF)

**S6 Fig.** Timeseries of climate variability index/probability (PDO, ONI, NPGO, and MHW; left y-axis) and sperm whale presence (black points; right y-axis) for CB. Sperm whale presence for the PDO, ONI, and NPGO were normalized between −1 and 1 to align with the respective climate variability index axis.

(TIFF)

**S7 Fig.** Timeseries of climate variability index/probability (PDO, ONI, NPGO, and MHW; left y-axis) and sperm whale presence (black points; right y-axis) for GOA. Sperm whale presence for the PDO, ONI, and NPGO were normalized between −1 and 1 to align with the respective climate variability index axis.

(TIFF)

**S8 Fig.** Timeseries of climate variability index/probability (PDO, ONI, NPGO, and MHW; left y-axis) and sperm whale presence (black points; right y-axis) for All-Sites. Sperm whale presence for the PDO, ONI, and NPGO were normalized between −1 and 1 to align with the respective climate variability index axis.

(TIFF)

**S9 Fig.** GLM plots displaying the relationship between mean monthly presence of sperm whales for site CB and the PDO, ONI, and NPGO index. All PDO and NPGO plots represent an eight-month lag, except for the Social Groups which does not include a lag. All ONI plots represent a nine-month lag. Each row (and color) represents outputs from the different size class models for each variable: Inclusive, Social Groups, Mid-Size, and Adult Males. All plots include 95% confidence intervals represented by the grey shading surrounding the linear regression. The regression formula for each model is displayed in the top left-hand corner. Covariates that were not retained in the model or not significant are represented with ‘NA’.

(TIFF)

**S10 Fig.** GLM plots displaying the relationship between mean monthly presence of sperm whales for the GOA region and the PDO, ONI, and NPGO index. All PDO and NPGO plots represent an eight-month lag, and all ONI plots represent a nine-month lag. Each row (and color) represents outputs from the different size class models for each variable: Inclusive, Social Groups, Mid-Size, and Adult Males. All plots include 95% confidence intervals represented by the grey shading surrounding the linear regression. The regression formula for each model is displayed in the top left-hand corner. Covariates that were not retained in the model or not significant are represented with ‘NA’.

(TIFF)

**S2 Table.** Generalized linear model (GLM) summaries testing the relationship of sperm whale presence and the Pacific Decadal Oscillation (PDO), Oceanic Nino Index (ONI), North Pacific Gyre Oscillation (NPGO), and Marine Heat Wave (MHW) indices for all GAM/GEE models that included year as a variable (i.e., greater than 5 years of data). For each model and class, significance of the model with no lag is denoted with an asterisk (*) in the column ‘Sig’. Significance of a model with a lag is denoted by a value from 8 to 12 representing the number of lags in months in the column ‘Sig’. Respective p-values and R^2^ values for each GLM is denoted for each significant model. Models that were not significant are denoted by ‘NA’. Models where year was not significant in the corresponding GAM/GEE model are *italicized*.

(DOCX)

## Acknowledgements

The authors would like to acknowledge the following agencies for funding and support during this study: U.S. Pacific Fleet, specifically Chip Johnson, Christiana Salles, and Jessica Bredvik, Pacific Life Foundation, specifically Bob Haskell. Support was also provided by the U.S. Fish and Wildlife Service crew with the R/V Tiglax for instrument deployment and recovery. We thank Bruce Thayre, John Hurwitz, and Ryan Griswold for coordinating instrument deployments and recoveries and Erin O’Neill for data processing. We also thank Tiago Marques for providing constructive feedback to improve the manuscript.

## Author Contributions

**Conceptualization:** Natalie Posdaljian, Alba Solsona-Berga, John A. Hildebrand, Simone Baumann-Pickering

**Data curation:** Natalie Posdaljian, Sean M. Wiggins, Kieran Lenssen

**Formal analysis:** Natalie Posdaljian, Caroline Soderstjerna

**Funding acquisition:** John A. Hildebrand, Simone Baumann-Pickering

**Investigation:** Natalie Posdaljian, Simone Baumann-Pickering

**Methodology:** Natalie Posdaljian, Alba Solsona-Berga, Sean M. Wiggins, Kieran Lenssen, Simone Baumann-Pickering

**Project administration:** Simone Baumann-Pickering

**Software:** Natalie Posdaljian, Alba Solsona-Berga, John A. Hildebrand, Sean M. Wiggins

**Supervision:** John A. Hildebrand, Simone Baumann-Pickering

**Validation:** Natalie Posdaljian

**Visualization:** Natalie Posdaljian

**Writing – original draft:** Natalie Posdaljian

**Writing – review & editing:** Alba Solsona-Berga, John A. Hildebrand, Caroline Soderstjerna, Sean M. Wiggins, Kieran Lenssen, Simone Baumann-Pickering

## References

1. Rice DW. Sperm whale Physter macrocephalus Linneaus. In: Ridgway SH, Harrison RJ, editors. Handbook of Marine Mammals. London, U.K.: Academic Press; 1989. p. 177–233.

2. Best PB. Social organization in sperm whales, Physeter macrocephalus. In: Behavior of marine animals. Springer; 1979. p. 227–89.

3. Gregr EJ, Trites AW. Predictions of critical habitat for five whale species in the waters of coastal British Columbia. Can J Fish Aquat Sci. 2001;58(7):1265–85.

4. Whitehead H. Society and culture in the deep and open ocean: The sperm whale and other cetaceans. In: Animal social complexity: Intelligence, culture, and individualized societies. Harvard University Press; 2003.

5. Carretta JV, Forney KA, Oleson EM, Weller DW, Lang AR, Baker J, et al. U.S. Pacific Marine Mammal Stock Assessments: 2019. 2020;(August).

6. Mesnick SL, Taylor BL, Archer FI, Martien KK, Treviño SE, Hancock-Hanser BL, et al. Sperm whale population structure in the eastern and central North Pacific inferred by the use of single-nucleotide polymorphisms, microsatellites and mitochondrial DNA. Mol Ecol Resour. 2011;11(SUPPL. 1):278–98.

7. Tomlin AG. Mammals of the USSR and Adjacent Countries. Cetacea. Israel Program Sci. Transl. 9 (124): 717. Natl Tech Inf Serv TT. 1967;65–50086.

8. Mizroch SA, Rice DW. Ocean nomads: Distribution and movements of sperm whales in the North Pacific shown by whaling data and Discovery marks. Mar Mammal Sci. 2013;29(2):136–65.

9. Nishiwaki M. Distribution and migration of the larger cetaceans in the North Pacific as shown by Japanese whaling results. Whales, dolphins porpoises Univ Calif Press Berkeley. 1966;171–91.

10. Berzin AA. Kashalot (The sperm whale). Moscow: Pischevaya Promyshlenmost; 1972.

11. Fearnbach H, Durban JW, Mizroch SA, Barbeaux S, Wade PR. Winter observations of a group of female and immature sperm whales in the high-latitude waters near the Aleutian Islands, Alaska. Mar Biodivers Rec. 2014;5(3):1–4.

12. Ivashchenko YV., Brownell RL, Clapham PJ. Distribution of Soviet catches of sperm whales Physeter macrocephalus in the North Pacific. Endanger Species Res. 2014;25(3):249–63.

13. Berzin AA, Rovnin AA. The distribution and migrations of whales in the northeastern part of the Pacific, Chukchi, and Bering Seas. Iszv Tikhookean Nauchno Issled Inst Ryb Khoz Okeanogr. 1966;58:179–208.

14. Berzin AA. The Sperm Whale, Israel Program for Scientific Translation, Jerusalem. Transl from Russ Berzin. 1971;

15. Rone BK, Zerbini AN, Douglas AB, Weller DW, Clapham PJ. Abundance and distribution of cetaceans in the Gulf of Alaska. Mar Biol. 2017;164(1):1–23.

16. Watwood SL, Miller PJO, Johnson M, Madsen PT, Tyack PL. Deep-diving foraging behaviour of sperm whales (Physeter macrocephalus). J Anim Ecol. 2006;75(3):814–25.

17. Watkins WA. Acoustics and the behavior of sperm whales. In: Animal Sonar Systems. Springer; 1980. p. 283–90.

18. Mellinger K, Stafford M, Fox G. Seasonal Occurence of Sperm Whale (Physeter Macrocephalus) Sounds in the Gulf of Alaska, 1999-2001. Mar Mammal Sci. 2004;20(January):48–62.

19. Diogou N, Palacios DM, Nieukirk SL, Nystuen JA, Papathanassiou E, Katsanevakis S, et al. Sperm whale (Physeter macrocephalus) acoustic ecology at Ocean Station PAPA in the Gulf of Alaska – Part 1: Detectability and seasonality. Deep Res Part I Oceanogr Res Pap [Internet]. 2019;150(August 2018):103047. Available from: https://doi.org/10.1016/j.dsr.2019.05.007

20. Rice A, Širović A, Trickey JS, Debich AJ, Gottlieb RS, Wiggins SM, et al. Cetacean occurrence in the Gulf of Alaska from long-term passive acoustic monitoring. Mar Biol [Internet]. 2021;1–29. Available from: https://doi.org/10.1007/s00227-021-03884-1

21. Gordon JCD. Evaluation of a method for determining the length of sperm whales (*Physeter catodon*) from their vocalizations. J Zool London. 1991;224:301–14.

22. Growcott A, Miller B, Sirguey P, Slooten E, Dawson S. Measuring body length of male sperm whales from their clicks: The relationship between inter-pulse intervals and photogrammetrically measured lengths. J Acoust Soc Am. 2011;130(1):568–73.

23. Solsona-Berga A, Posdaljian N, Hildebrand JA, Baumann-Pickering S. Echolocation repetition rate as a proxy to monitor population structure and dynamics of sperm whales. Remote Sens Ecol Conserv. 2022;8(6):827–40.

24. Møhl B, Wahlberg M, Madsen PT, Miller LA, Surlykke A. Sperm whale clicks: Directionality and source level revisited. J Acoust Soc Am. 2000;107(1):638–48.

25. Barlow J, Taylor BL. Estimates of Sperm Whale Abundance in the Northeastern Temperate Pacific From a Combined Acoustic and Visual Survey. Mar Mammal Sci. 2005;21(3):429–45.

26. Madsen PT, Wahlberg M, Møhl B. Male sperm whale (Physeter macrocephalus) acoustics in a high-latitude habitat: Implications for echolocation and communication. Behav Ecol Sociobiol. 2002;53(1):31–41.

27. Mathias D, Thode AM, Straley J, Andrews RD. Acoustic tracking of sperm whales in the Gulf of Alaska using a two-element vertical array and tags. J Acoust Soc Am. 2013;134(3):2446–61.

28. Backus RH, Schevill WE. Whales, dolphins and porpoises. University of California Press; 1966. 510–528 p.

29. Norris KS, Harvey GW. A Theory for the Function of the Spermaceti Organ of the Sperm Whale (Physeter catodon L.). Anim Obs Navig. 1972;397–417.

30. Nishiwaki M, Ohsumi S, Maeda Y. Change of form in the sperm whale accompanied with growth. Sci Reports Whales Res Inst. 1963;17:1–14.

31. Caruso F, Sciacca V, Bellia G, Domenico E De, Larosa G, Papale E, et al. Size Distribution of Sperm Whales Acoustically Identified during Long Term Deep-Sea Monitoring in the Ionian Sea Size Distribution of Sperm Whales Acoustically Identified during Long Term Deep-Sea Monitoring in the Ionian Sea. PLoS One. 2015;1–16.

32. Beslin WAM, Whitehead H, Gero S, Beslin WAM, Whitehead H. Automatic acoustic estimation of sperm whale size distributions achieved through machine recognition of on-axis clicks. J Acoust Soc Am. 2018;144(6):3485–95.

33. Møhl B, Wahlberg M, Madsen PT, Heerfordt A, Lund A. The monopulsed nature of sperm whale clicks. J Acoust Soc Am. 2003;114(2):1143–54.

34. Zimmer WMX, Madsen PT, Teloni V, Tyack MPJ and PL. Off-axis effects on the multipulse structure of sperm whale usual clicks with implications for sound production. J Acoust Soc Am 118(5)3337-3345 2005. 2005;118(November):9.

35. Goold JC, Jones SE. Time and frequency domain characteristics of sperm whale clicks. J Acoust Soc Am. 1995;98(3):1279–91.

36. Jensen FH, Johnson M, Ladegaard M, Wisniewska DM, Madsen PT. Narrow Acoustic Field of View Drives Frequency Scaling in Toothed Whale Biosonar. Curr Biol [Internet]. 2018;1–8. Available from: https://www.sciencedirect.com/science/article/pii/S0960982218313666?via%3Dihub

37. Wiggins SM, Hildebrand JA. High-frequency Acoustic Recording Package (HARP). Int Symp Underw Technol 2007 Int Work Sci Use Submar Cables Relat Technol 2007. 2007;(April):17–20.

38. Solsona-Berga A, Frasier KE, Baumann-Pickering S, Hildebrand JA. DetEdit, a Graphical User Interface for Annotating and Editing Events Detected in Acoustic Data. PLOS Comput Biol. 2020;1–10.

39. Inc. M. MATLAB Version: 9.1.0 (2016b) [Internet]. Natick, Massachusetts: The MathWorks Inc. 2016. Available from: https://www.mathworks.com

40. Welch PD. The use of fast Fourier transform for the estimation of power spectra: A method based on time averaging over short, modified periodograms. 1967;(2):70–3.

41. Jaquet N. A simple photogrammetric technique to measure sperm whales at sea. Mar Mammal Sci. 2006 Oct;22(4):862–79.

42. Jochens a., Jochens a., Biggs D, Biggs D, Benoit-Bird K, Benoit-Bird K, et al. Sperm whale seismic study in the Gulf of Mexico. Synthesis Report. Mms [Internet]. 2008;96(5):3268. Available from: http://link.aip.org/link/JASMAN/v96/i5/p3268/s3&Agg=doi

43. Hastie T, Tibshirani R. Generalized additive models: Some applications. J Am Stat Assoc. 1987;82(398):371–86.

44. Liang KY, Zeger SL. Longitudinal data analysis using generalized linear models. Biometrika. 1986;73(1):13–22.

45. Pirotta E, Matthiopoulos J, Mackenzie M, Scott-hayward L, Rendell L. Modelling sperm whale habitat preference : a novel approach combining transect and follow data. Mar Ecol Prog Ser. 2011;436:257–72.

46. R Core Team. R: A language and environment for statistical computing. R Found Stat Comput [Internet]. 2022; Available from: https://www.r-project.org/

47. Panigada S, Zanardelli M, MacKenzie M, Donovan C, Mélin F, Hammond PS. Modelling habitat preferences for fin whales and striped dolphins in the Pelagos Sanctuary (Western Mediterranean Sea) with physiographic and remote sensing variables. Remote Sens Environ. 2008;112(8):3400–12.

48. Benjamins S, van Geel N, Hastie G, Elliott J, Wilson B. Harbour porpoise distribution can vary at small spatiotemporal scales in energetic habitats. Deep Res Part II Top Stud Oceanogr [Internet]. 2017;141:191–202. Available from: http://dx.doi.org/10.1016/j.dsr2.2016.07.002

49. Merkens K, Simonis A, Oleson E. Geographic and temporal patterns in the acoustic detection of sperm whales Physeter macrocephalus in the central and western North Pacific Ocean. Endanger Species Res. 2019;39:115–33.

50. Bailey H, Corkrey R, Cheney B, Thompson PM. Analyzing temporally correlated dolphin sightings data using generalized estimating equations. Mar Mammal Sci. 2013;29(1):123–41.

51. Booth CG, Embling C, Gordon J, Calderan S V., Hammond PS. Habitat preferences and distribution of the harbour porpoise Phocoena phocoena west of Scotland. Mar Ecol Prog Ser. 2013;478:273–85.

52. Stimpert AK, DeRuiter SL, Falcone EA, Joseph J, Douglas AB, Moretti DJ, et al. Sound production and associated behavior of tagged fin whales (Balaenoptera physalus) in the Southern California Bight. Anim Biotelemetry. 2015;3(1):1–12.

53. Zuur AF. A beginner’s guide to generalized additive models with R (pp. 1-206). Mater Sci Eng A [Internet]. 2012;27(1):1–14. Available from: https://www.tandfonline.com/doi/full/10.1080/02670836.2016.1231746%0Ahttp://dx.doi.org/10.1016/j.actamat.2011.03.055%0Ahttp://dx.doi.org/10.1016/j.msea.2016.02.076%0A http://dx.doi.org/10.1016/j.msea.2012.06.095%0Ahttps://doi.org/10.1016/j.ijhydene.2019.11

54. Fox J. John Fox and Sanford Weisberg. An R companion to Appl regression, 3rd ed Sage Publ Sch. 2019;

55. Halekoh U, Højsgaard S, Yan J. The R Package geepack for Generalized Estimating Equations [Internet]. Vol. 15, JSS Journal of Statistical Software. 2006. Available from: http://www.jstatsoft.org/

56. Wood SN. Fast stable restricted maximum likelihood and marginal likelihood estimation of semiparametric generalized linear models. J R Stat Soc Ser B Stat Methodol. 2011;73(1):3–36.

57. Pan W. Akaike’s Informat ion Criterion in Generalized Estimating Equations. Biometrics. 2001;57(March):120–5.

58. Halekoh U, Højsgaard S, Yan J. The R package geepack for generalized estimating equations. J Stat Softw. 2006;15(2):1–11.

59. Lüdecke D, Ben-Shachar M, Patil I, Waggoner P, Makowski D. performance: An R Package for Assessment, Comparison and Testing of Statistical Models. J Open Source Softw. 2021 Apr 21;6(60):3139.

60. Tjur T. Coefficients of determination in logistic regression models - A new proposal: The coefficient of discrimination. Am Stat [Internet]. 2009;63(4):366–72. Available from: https://doi.org/10.1198/tast.2009.08210

61. Gelman A, Hill J. Data analysis using regression and multilevel/hierarchical models. Cambridge University Press; 2007. 625 p.

62. Albers S. Rsoi: Import Various Northern and Southern Hemisphere Climate Indices [Internet]. 2020. Available from: https://cran.r-project.org/package=rsoi

63. Jacox MG, Alexander MA, Amaya D, Becker E, Bograd SJ, Brodie S, et al. Global seasonal forecasts of marine heatwaves. Nature. 2022;604(7906):486–90.

64. Rob J. Hyndman, Yeasmin Khandakar. Automatic Time Series Forecasting: The forecast Package for R. J Stat Softw [Internet]. 2008;27(3):22. Available from: http://www.jstatsoft.org/%0Ahttp://www.jstatsoft.org/v27/i03/paper

65. Hyndman R, Athanasopoulos G, Bergmeir C, Caceres G, Chhay L, O’Hara-Wild M, Petropoulos F, et al. forecast: Forecasting functions for time series and linear models [Internet]. R package version 8.21. 2023. Available from: https://pkg.robjhyndman.com/forecast/

66. Diogou N, Palacios DM, Nystuen JA, Papathanassiou E, Katsanevakis S, Klinck H. Sperm whale (Physeter macrocephalus)acoustic ecology at Ocean Station PAPA in the Gulf of Alaska – Part 2: Oceanographic drivers of interannual variability. Deep Res Part I Oceanogr Res Pap. 2019;150(August 2018):103044.

67. Di Lorenzo E, Schneider N, Cobb KM, Franks PJS, Chhak K, Miller AJ, et al. North Pacific Gyre Oscillation links ocean climate and ecosystem change. Geophys Res Lett. 2008;35(8):2–7.

68. Omura H. On the Body. Weight of Sperm and Sei Whales located in the Adjacent Waters of Japan. Sci Reports Whales Res Inst. 1950;4:27–113.

69. Dufault S, Whitehead H, Dillon M. An examination of the current knowledge on the stock structure of sperm whales (Physeter macrocephalus) worldwide. J Cetacean Res Manag. 1999;1(1):1–10.

70. Nowak RM. Walker’s Marine Mammals of the World. Baltimore: The Johns Hopkins University Press; 2003.

71. McClain CR, Balk MA, Benfield MC, Branch TA, Chen C, Cosgrove J, et al. Sizing ocean giants: Patterns of intraspecific size variation in marine megafauna. PeerJ. 2015;2015(1).

72. Gaskin DE, MW C. Sperm whales (Physeter catodon L.) in the Cook Strait region of New Zealand: some data on age, growth and mortality. 1973;

73. Gaskin DE. Composition of schools of sperm whales physeter catodon linn. East of New Zealand. New Zeal J Mar Freshw Res. 1970;4(4):456–71.

74. Hamer DJ, Childerhouse SJ, Gales NJ. Odontocete bycatch and depredation in longline fisheries: A review of available literature and of potential solutions. Mar Mammal Sci. 2012;28(4):345–74.

75. Sigler MF, Lunsford CR, Straley JM, Liddle JB. Sperm whale depredation of sablefish longline gear in the northeast Pacific Ocean. Mar Mammal Sci. 2008;24(1):16–27.

76. Kasuya T, Miyashita T. Distribution of sperm whale stocks in the North Pacific. Sci Reports Whales Res Inst Tokyo 3931-75 1988. 1988;39:45.

77. Straley JM, Schorr GS, Thode AM, Calambokidis J, Lunsford CR, Chenoweth EM, et al. Depredating sperm whales in the Gulf of Alaska: Local habitat use and long distance movements across putative population boundaries. Endanger Species Res. 2014;24(2):125–35.

78. Mullin KD, Steiner L, Dunn C, Claridge D, García LG, Gordon J, et al. Long-Range Longitudinal Movements of Sperm Whales (Physeter macrocephalus) in the North Atlantic Ocean Revealed by Photo-Identification. Aquat Mamm. 2022;48(1):3–8.

79. Jaquet N, Gendron D. Distribution and relative abundance of sperm whales in relation to key environmental features, squid landings and the distribution of other cetacean species in the Gulf of California, Mexico. Mar Biol. 2002;141(3):591–601.

80. Whitehead H, Arnbom T. Social organization of sperm whales off the Galapagos Islands, February–April 1985. Can J Zool [Internet]. 1987;65(4):913–9. Available from: http://www.nrcresearchpress.com/doi/10.1139/z87-145

81. Kato H. Observations on tooth scars on the head of male sperm whale, as an indication of intra-sexual fightings. Sci Reports Whales Res Inst. 1984;35(35):39–46.

82. Whitehead H. The behaviour of mature male sperm whales on the Galapagos Islands breeding grounds. Can J Zool. 1993;71(4):689–99.

83. Gero S, Gordon JCD, Carlson C, Evans P, Whitehead H. Population estimate and inter-island movement of sperm whales, *Physeter macrocephalus*, in the Eastern Caribbean. Vol. 9, Journal of Cetacean Research and Management. 2007. p. 143–50.

84. Vachon F, Hersh TA, Rendell L, Gero S, Whitehead H. Ocean nomads or island specialists? Culturally driven habitat partitioning contrasts in scale between geographically isolated sperm whale populations. R Soc Open Sci. 2022;9(5).

85. Engelhaupt D, Rus Hoelzel A, Nicholson C, Frantzis A, Mesnick S, Gero S, et al. Female philopatry in coastal basins and male dispersion across the North Atlantic in a highly mobile marine species, the sperm whale (Physeter macrocephalus). Mol Ecol. 2009;18(20):4193–205.

86. Carpinelli E, Gauffier P, Verborgh P, Airoldi S, David L, Di-Méglio N, et al. Assessing sperm whale (Physeter macrocephalus) movements within the western Mediterranean Sea through photo-identification. Aquat Conserv Mar Freshw Ecosyst. 2014;24(SUPPL.1):23– 30.

87. Whitehead H, Christal J, Dufault S. Past and distant whaling and the rapid decline of sperm whales off the Galapagos Islands. Conserv Biol. 1997;11(6):1387–96.

88. Jaquet N, Whitehead H. Scale-dependent correlation of sperm whale distribution with environmental features and productivity in the South Pacific. Mar Ecol Prog Ser. 1996;135(1–3):1–9.

89. Zador S. Appendix C Ecosystem considerations for 2012 [Internet]. 2011. Available from: http://www.afsc.noaa.gov/REFM/docs/2011/ecosystem.pdf

90. Boyd PW. The NE subarctic Pacific in winter: I. Biological standing stocks. Mar Ecol Prog Ser. 1995;128(1–3):11–24.

91. Whitney FA, Freeland HJ. Variability in upper-ocean water properties in the NE Pacific Ocean. Deep Res Part II Top Stud Oceanogr. 1999;46(11–12):2351–70.

92. Morato T, Varkey DA, Damaso C, Machete M, Santos M, Prieto R, et al. Evidence of a seamount effect on aggregating visitors. Mar Ecol Prog Ser. 2008;357(Fonteneau 1991):23–32.

93. Wong SNP, Whitehead H. Seasonal occurrence of sperm whales (Physeter macrocephalus) around Kelvin Seamount in the Sargasso Sea in relation to oceanographic processes. Deep Res Part I Oceanogr Res Pap. 2014;

94. Dede A, Öztürk AA, Tonay AM, Uğur Ö, Gönülal O, Öztürk B, et al. Cetacean sightings in the Finike Seamounts area and adjacent waters during the surveys in 2021. J Black Sea Mediterr Environ. 2022;28(2):221–39.

95. Gregr EJ, Nichol L, Ford JKB, Ellis G, Trites AW. Migration and population structure of northeastern Pacific whales off coastal British Columbia: An analysis of commercial whaling records from 1908-1967. Mar Mammal Sci. 2000;16(4):699–727.

96. Mizroch SA, Rice DW. Have North Pacific killer whales switched prey species in response to depletion of the great whale populations? Mar Ecol Prog Ser. 2006;310:235–46.

97. Pike GC. Schooling and migration of sperm whales off British Columbia. Unpubl Manuscript, 18pp. 1965;

98. Ohsumi S. Reproduction of the sperm whale in the north-west Pacific. Sci Reports Whales Res Institute, Tokyo. 1965;19:1–35.

99. Okutani T, Nemoto T. Squids as the food of sperm whales in the Bering Sea and the Alaska Gulf. Tokai Reg Fish Lab Sci Rep Whales Res Inst [Internet]. 1964;18. Available from: http://www.icrwhale.org/pdf/SC018111-122.pdf

100. Santos MB, Pierce GJ, Boyle PR, Reid RJ, Ross HM, Patterson IAP, et al. Stomach contents of sperm whales Physeter macrocephalus stranded in the North Sea 1990-1996. Mar Ecol Prog Ser. 1999;183(June):281–94.

101. Fristrup KM, Harbison GR. How do sperm whales catch squids? Mar Mammal Sci. 2002;18(1):42–54.

102. Das K, Lepoint G, Leroy Y, Bouquegneau JM. Marine mammals from the southern North Sea: Feeding ecology data from δ13C and δ15N measurements. Mar Ecol Prog Ser. 2003;263(January):287–98.

103. Wild LA, Mueter F, Witteveen B, Straley JM. Exploring variability in the diet of depredating sperm whales in the Gulf of Alaska through stable isotope analysis. R Soc Open Sci. 2020;7(3).

104. Ormseth OA. Assessment of the squid stock complex in the Gulf of Alaska. Executive Summary. 2017.

105. Gregr EJ, Baumgartner MF, Laidre KL, Palacios DM. Marine mammal habitat models come of age: The emergence of ecological and management relevance. Endanger Species Res. 2013;22(3):205–12.

106. Palacios DM, Baumgartner MF, Laidre KL, Gregr EJ. Beyond correlation: Integrating environmentally and behaviourally mediated processes in models of marine mammal distributions. Endanger Species Res. 2013;22(3):191–203.

107. Mantua NJ, Hare SR, Zhang Y, Wallace JM, Francis RC. A Pacific Interdecadal Climate Oscillation with Impacts on Salmon Production. Bull Am Meteorol Soc. 1997;78(6):1069–79.

108. Anderson PJ, Piatt JF. Community reorganization in the Gulf of Alaska following ocean climate regime shift. Mar Ecol Prog Ser. 1999;189:117–23.

109. Brodeur RD, Ware DM. Long-term variability in zooplankton biomass in the subarctic Pacific Ocean. Fish Oceanogr. 1992;1(1):32–8.

110. Hatch SA. Kittiwake diets and chick production signal a 2008 regime shift in the Northeast Pacific. Mar Ecol Prog Ser. 2013;477:271–84.

111. Anderson PJ. Declines of forage species in the Gulf of Alaska, 1972-1995, as an indicator of regime shift. In: Forage fishes in marine ecosystems. University of Alaska; 1997.

112. Piatt JF, Anderson P. Response of Common Murres to the Exxon Valdez Oil Spill and Long-Term Changes in the Gulf of Alaska Marine Ecosystem. Am Fish Soc Symp. 1996;18:720–37.

113. Merrick RL, Chumbley MK, Byrd GV. Diet diversity of Steller sea lions (Eumetopias jubatus) and their population decline in Alaska: a potential relationship. Can J Fish Aquat Sci. 1997;54(6):1342–8.

114. Rodhouse PG, Hatfield EMC. Age determination in squid using statolith growth increments.pdf. Fish Res. 1990;8:323–34.

115. Forsythe JW. Accounting for the effect of temperature on squid growth in nature: From hypothesis to practice. Mar Freshw Res. 2004;55(4):331–9.

116. Crawford W, Mckinnell S. Continuing Cool in the Northeast Pacific Ocean. 2013;21(1):2012–3.

117. Stewart JS, Hazen EL, Bograd SJ, Byrnes JEK, Foley DG, Gilly WF, et al. Combined climate- and prey-mediated range expansion of Humboldt squid (Dosidicus gigas), a large marine predator in the California Current System. Glob Chang Biol. 2014;20(6):1832–43.

118. Peterson WT, Robert M, Bond N. The Blob is gone but has morphed into a strongly positive PDO/SST pattern. PICES Press. 2016;21(2):44–6.

119. Crawford, W.; McKinnell, S.; and Freeland H. News of the Northeast Pacific Ocean. PICES Press [Internet]. 2012;20(1):1–3. Available from: http://search.proquest.com/openview/1cfdcfd44cfd542280c90aefbe6718ca/1?pq-origsite=gscholar&cbl=666306

120. Jackson JM, Myers PG, Ianson D. An examination of advection in the northeast Pacific Ocean, 2001-2005. Geophys Res Lett. 2006;33(15):2001–5.

121. NOAA Climate Prediction Center. Oceanic Niño Index [Internet]. 2023. Available from: https://origin.cpc.ncep.noaa.gov/products/analysis_monitoring/ensostuff/ONI_v5.php

122. Whitney FA, Welch DW. Impact of the 1997-1998 El Niño and 1999 La Niña on nutrient supply in the Gulf of Alaska. Prog Oceanogr. 2002;54(1–4):405–21.

123. Mantua NJ, Hare SR. The Pacific Decadal Oscillation. Vol. 58, Journal of Oceanography. 2002. p. 35–44.

124. Gershunov A, Barnett TP. Interdecadal Modulation of ENSO Teleconnections. Bull Am Meteorol Soc. 1998;79(12):2715–25.

125. Cummins PF, Lagerloef GSE. Low-frequency pycnocline depth variability at Ocean Weather Station P in the northeast Pacific. J Phys Oceanogr. 2002;32(11):3207–15.

126. Mantua NJ. Pacific Decadal Oscillation (PDO) [Internet]. 2023 [cited 2023 Oct 1]. Available from: https://www.ncei.noaa.gov/access/monitoring/pdo/

127. Litzow MA, Ciannelli L, Puerta P, Wettstein JJ, Rykaczewski RR, Opiekun M. Non-stationary climate-salmon relationships in the Gulf of Alaska. Proc R Soc B Biol Sci. 2018;285(1890).

128. Bond NA, Cronin MF, Freeland H, Mantua N. Causes and impacts of the 2014 warm anomaly in the NE Pacific. Geophys Res Lett. 2015;42(9):3414–20.

129. Yang B, Emerson SR, Angelica Penã M. The effect of the 2013-2016 high temperature anomaly in the subarctic Northeast Pacific (the “blob”) on net community production. Biogeosciences. 2018;15(21):6747–59.

130. Cavole LM, Demko AM, Diner RE, Giddings A, Koester I, Pagniello CMLS, et al. Biological impacts of the 2013–2015 warm-water anomaly in the northeast Pacific: Winners, Losers, and the Future. Oceanography. 2016;29(2):273–85.

131. Hazen EL, Jorgensen S, Rykaczewski RR, Bograd SJ, Foley DG, Jonsen ID, et al. Predicted habitat shifts of Pacific top predators in a changing climate. Nat Clim Chang. 2013 Mar;3(3):234–8.

132. Baumann-Pickering S, Trickey JS, Wiggins SM, Oleson EM. Odontocete occurrence in relation to changes in oceanography at a remote equatorial Pacific seamount. Mar Mammal Sci. 2016;32(3):805–25.

133. Griffin RB. Sperm whale distributions and community ecology associated with a warm-core ring off Georges Bank. Mar Mammal Sci. 1999;15(1):33–51.

